# ARL13B-Cerulean rescues *Arl13b*-null mouse from embryonic lethality and reveals a role for ARL13B in spermatogenesis

**DOI:** 10.1101/2025.03.24.644968

**Authors:** Alyssa B. Long, Isabella M. Wilson, Tiffany T. Terry, Robert E. Van Sciver, Tamara Caspary

## Abstract

ARL13B is a regulatory GTPase enriched in cilia, making it a popular marker for this organelle. *Arl13b^hnn/hnn^* mice lack ARL13B expression, die during midgestation, and exhibit defects in ciliogenesis. The *R26Arl13b-Fucci2aR* biosensor mouse line directs the expression of fluorescently tagged full-length *Arl13b* cDNA upon *Cre* recombination. To determine whether constitutive, ubiquitous expression of ARL13B-Cerulean can replace endogenous gene expression, we generated *Arl13b^hnn/hnn^* animals expressing ARL13B-Cerulean. We show that *Arl13b^hnn/hnn^;Arl13b-Cerulean* mice survive to adulthood with no obvious physical or behavioral defects, indicating that the fluorescently tagged protein can functionally replace the endogenous protein during development. However, we observed that rescued males failed to sire offspring, revealing a role for ARL13B in spermatogenesis. This work shows that the *R26Arl13b-Fucci2aR* mouse contains an inducible allele of *Arl13b* capable of functioning in most tissues and biological processes.

## INTRODUCTION

ARL13B is a regulatory GTPase that is enriched in cilia and is required for proper ciliogenesis and trafficking of signaling molecules (Caspary et al., 2007; Higginbotham et al., 2012; Larkins et al., 2011; Sun et al., 2004). Cilia are microtubule-based projections that can be non-motile or motile. Several transgenic models label cilia by fusing ARL13B to fluorescent proteins (Bangs et al., 2015; Borovina et al., 2010; Delling et al., 2013; Ford et al., 2018; Schmitz et al., 2017). These models display no major developmental defects; embryos become viable adults and appear healthy overall. To date, analyses of these fluorescently labeled ARL13B models are in wild-type animals, meaning solely in the context of ARL13B expressed from the endogenous locus.

Mice that are homozygous for an N-ethyl-N-nitrosourea-induced mutation, *Arl13b^hnn^*, die around embryonic day (E)13.5 and exhibit short cilia and defects in the structure of the ciliary axoneme (Caspary et al., 2007). The *Arl13b^hnn^* mutation disrupts the splicing of exon 2, resulting in the absence of ARL13B protein (Caspary et al., 2007). *Arl13b^hnn/hnn^* embryos display abnormal Hedgehog (Hh) signaling, consistent with cilia being required to transduce Hh signaling in vertebrates (Caspary et al., 2007; Huangfu et al., 2003). Indeed, components of vertebrate Hh signaling are abnormally trafficked in *Arl13b^hnn^* cilia, wherein the obligate pathway transducer Smoothened is localized to cilia in the absence of stimulation (Larkins et al., 2011).

Conditional alleles of *Arl13b* circumvent embryonic lethality and show that ARL13B is critical for a variety of developmental processes. In the kidney, deletion of *Arl13b* leads to cysts in mice and zebrafish (Augière et al., 2024; Bay et al., 2018; Duldulao et al., 2009; Li et al., 2016; Seixas et al., 2016; Sun et al., 2004). *Arl13b* deletion throughout the murine central nervous system leads to hydrocephaly (Su et al., 2012; Suciu et al., 2021). *Arl13b* deletion in specific subsets of neurons or interneurons reveals brain defects in mice including small cerebellar vermis, impaired interneuron migration, and abnormal radial glial scaffolding (Guo et al., 2017; Higginbotham et al., 2012; Higginbotham et al., 2013; Suciu et al., 2021). ARL13B also functions in the specialized sensory cilia of the visual and olfactory systems (Dilan et al., 2019; Fiore et al., 2020; Habif et al., 2023; Hanke-Gogokhia et al., 2017; Joiner et al., 2015). Patients with the ciliopathy Joubert syndrome (JS) exhibit similar phenotypes, consistent with *ARL13B* mutations causing JS (OMIM 612291) (Cantagrel et al., 2008; Thomas et al., 2015).

*Arl13b^V358A/V358A^* mice are viable and fertile despite lacking ARL13B protein specifically in cilia (Gigante et al., 2020). This engineered protein variant, ARL13B^V358A^, retains the known biochemical activities of ARL13B but is not detectable in cilia (Gigante et al., 2020; Higginbotham et al., 2012; Mariani et al., 2016). *Arl13b^V358A/V358A^* mice display enlarged, cystic kidneys, develop hyperphagia, and become obese, indicating ciliary ARL13B functions to regulate kidney development as well as energy homeostasis (Terry et al., 2023; Van Sciver et al., 2023). Hh signaling is normal in *Arl13b^V358A/V358A^* mice while ciliogenesis is abnormal, showing the two processes can be uncoupled and ciliary ARL13B regulates ciliogenesis in certain tissues (Gigante et al., 2020).

ARL13B’s function in ciliogenesis varies by cell type. *Arl13b* loss in tissues including the left-right organizer, neural tube, and male efferent ducts leads to short cilia, albeit at normal frequencies (Augière et al., 2024; Caspary et al., 2007; Duldulao et al., 2009; Larkins et al., 2011; Su et al., 2012). In contrast, cultured fibroblast cells derived from *Arl13b^hnn/hnn^* embryos exhibit short cilia at lower frequencies (Larkins et al., 2011). In the kidney, *Arl13b* deletion results in a complete absence of cilia (Duldulao et al., 2009; Li et al., 2016; Seixas et al., 2016; Sun et al., 2004). The mechanism(s) through which ARL13B controls ciliogenesis in any cell type remains unclear.

ARL13B expression levels correlate with cilia length in several cell types. While ARL13B loss leads to short or absent cilia, ARL13B overexpression can cause ciliary elongation (Larkins et al., 2011; Lu et al., 2015; Pintado et al., 2017). For example, cultured primary wild-type or *Arl13b^hnn/hnn^* fibroblasts overexpressing *Arl13b* display longer cilia than untransfected cells (Larkins et al., 2011). Ciliary lengthening reflects tissue-specific functions of ARL13B in ciliogenesis. In the *R26Arl13b-Fucci2aR* biosensor mouse line, the expression of full-length, fluorescently tagged ARL13B-Cerulean results in longer cilia in cultured cells, as well as *in vivo* in the mesenchyme, kidney tubules, and liver bile ducts (Ford et al., 2018). Meanwhile, cilia of the nasal epithelium and brain ependyma remain the same length as in control mice (Ford et al., 2018). This suggests ARL13B-Cerulean is functional and differentially impacts ciliogenesis in distinct cell types.

The *R26Arl13b-Fucci2aR* biosensor carries a floxed-STOP cassette between the CAG promoter and the full-length *Arl13b* cDNA fused to Cerulean, providing spatial and temporal control of expression (Ford et al., 2018). It also constitutively expresses mVenus and mCherry fused to fragments of hGem and hCdt1, enabling simultaneous monitoring of cilia and cell cycle progression (Ford et al., 2018; Sakaue-Sawano et al., 2008). The inclusion of full-length *Arl13b* cDNA suggests functional protein is produced and implies the *R26Arl13b-Fucci2aR* biosensor might also serve as an inducible *Arl13b* allele. To test this, we asked whether the recombined allele expressing ubiquitous, constitutive ARL13B-Cerulean protein from the *Rosa26* locus could replace endogenous ARL13B in the *Arl13b^hnn/hnn^* null mouse model. We also investigated the localization and level of ARL13B-Cerulean expression in several tissues where ARL13B functions.

## RESULTS

### Systemic expression of ARL13B-Cerulean rescues embryonic lethality of *Arl13b^hnn/hnn^* null mice

To generate *Arl13b^hnn/hnn^;Arl13b-Cerulean* mice, we first crossed *R26Arl13b-Fucci2aR* mice with animals carrying *CMV-Cre* to indelibly activate biosensor expression in all tissues (Fig. 1A). For simplicity, we call the constitutively-on allele *Arl13b-Cerulean.* We bred *Arl13b-Cerulean* mice to animals carrying *Arl13b^hnn^*, the *Arl13b-*null allele, and intercrossed the *Arl13b^hnn/+^;Arl13b-Cerulean* progeny. We determined the survival rate of pups two weeks after birth and performed genotyping. *Arl13b^hnn/+^;Arl13b-Cerulean* intercrosses generated no viable *Arl13b^hnn/hnn^* mice, which die at midgestation (Fig. 1B) (Caspary et al., 2007). In contrast, we observed *Arl13b^hnn/hnn^;Arl13b-Cerulean* mice at close to the predicted frequencies (Fig. 1B). The rescued *Arl13b^hnn/hnn^;Arl13b-Cerulean* mice survived into adulthood and displayed no gross morphological or behavioral defects (Fig. 1C). While we analyzed heterozygous and homozygous *Arl13b-Cerulean* animals, we generally did not observe any phenotypic differences, so we report them together unless specifically stated otherwise. We conclude that ARL13B-Cerulean functionally compensates for the loss of endogenous ARL13B protein during embryonic development, indicating it is an inducible allele.

**Figure 1.**
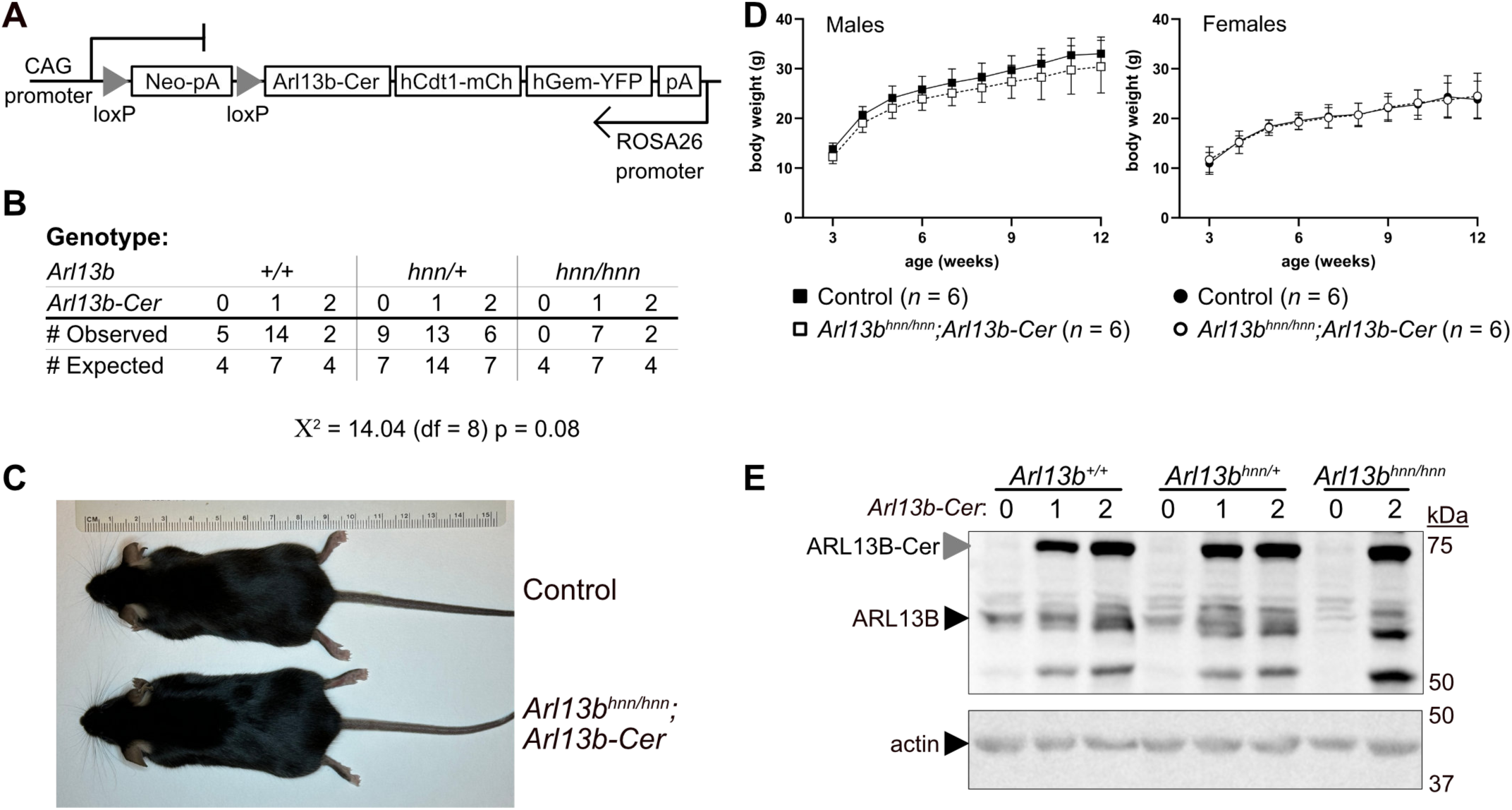
Arl13b^hnn/hnn^;Arl13b-Cerulean mice are viable. **A)** Schematic of the inducible *R26Arl13b-Fucci2aR* allele. **B)** Numbers of observed (*n* = 58) and expected pups from nine litters of *Arl13b^hnn/+^;Arl13b-Cerulean* intercrosses genotyped at 2 weeks. The genotype of endogenous *Arl13b* and the number of *Arl13b-Cerulean* alleles (0, 1, 2) are indicated. Chi-square test with 8 degrees of freedom (df) indicates the observed values do not significantly differ from expected; *p* value = 0.08. **C)** *Arl13b^hnn/+^;Arl13b-Cerulean* (control) and *Arl13b^hnn/hnn^;Arl13b-Cerulean* (rescue) female mice at 21 weeks. **D)** Weekly body weights of male and female control (black) and *Arl13b^hnn/hnn^;Arl13b-Cerulean* (white) mice, *n* = 6 for each group. Two-way ANOVA with Sidak’s multiple comparisons test was not significant at any time point for either sex; overall fixed effects (age by genotype) *p* values = 0.62 for males and 0.69 for females. **E)** Western blot of E12.5 whole-embryo lysates without (0) or with (1, 2 copies) *Arl13b-Cerulean*, probed with antibody against ARL13B (top image) or actin (bottom image).

As ciliary ARL13B regulates body weight, we investigated the growth of *Arl13b^hnn/hnn^;Arl13b-Cerulean* mice. We maintained cohorts of mice on breeder chow and measured body weight weekly from weaning (week three) to week twelve. We found no significant difference in weight curves between control (*Arl13b^+/+^* or *Arl13b^hnn/+^* with or without ARL13B-Cerulean) and *Arl13b^hnn/hnn^;Arl13b-Cerulean* male or female mice (Fig. 1D). *Arl13b^hnn/hnn^;Arl13b-Cerulean* mice maintained normal body weight throughout adulthood (Fig. S1A,B). These data indicate that ARL13B-Cerulean functions to regulate body weight homeostasis.

To evaluate the relative protein expression levels of ARL13B and ARL13B-Cerulean, we performed western blots using lysates from E12.5 embryos. We observed endogenous ARL13B at approximately 60 kDa in all embryos except the *Arl13b^hnn/hnn^* mutant (Fig. 1E). We detected the exogenous ARL13B-Cerulean fusion protein at approximately 72 kDa only in embryos carrying the biosensor (Fig. 1E). Lysates from embryos with two copies of the biosensor exhibited more ARL13B-Cerulean protein than those carrying one copy (Fig. 1E, S1C). In addition to protein bands at approximately 50 kDa in samples that express *Arl13b-Cerulean*, we also observed a band close to endogenous ARL13B’s size in lysate from the *Arl13b^hnn/hnn^;Arl13b-Cerulean* embryo (last lane of Fig. 1E), suggesting the fusion protein may be cleaved. The stronger signal produced by ARL13B-Cerulean compared to endogenous ARL13B likely reflected a high level of *Arl13b-Cerulean* expression driven by the CAG promoter, although we cannot rule out a higher affinity of the antibody to the fusion protein. Taken together, these results show that ARL13B-Cerulean is robustly expressed and rescues the embryonic lethality in *Arl13b^hnn/hnn^* animals.

### ARL13B-Cerulean expression restores wild-type levels of ciliation and cilia length to *Arl13b^hnn/hnn^* MEFs

To examine the ciliary localization of ARL13B and ARL13B-Cerulean in the *Arl13b^hnn/hnn^;Arl13b-Cerulean* mice, we isolated and immortalized mouse embryonic fibroblasts (MEFs) from wild-type, *Arl13b^hnn/hnn^*, and *Arl13b^hnn/hnn^;Arl13b-Cerulean* E12.5 embryos. We co-stained cilia with antibodies against glutamylated tubulin (GT335) and ARL13B. We detected both glutamylated tubulin and ARL13B in wild-type *Arl13b^+/+^* cilia (Fig. 2A). Mutant *Arl13b^hnn/hnn^* cells lacked ciliary ARL13B protein and displayed short cilia (Fig. 2B,E). We saw ARL13B-Cerulean localizing to cilia in *Arl13b^hnn/hnn^;Arl13b-Cerulean* cells using antibodies against either GFP, which recognizes the Cerulean tag, or ARL13B (Fig. 2C). These results show that ARL13B-Cerulean displays normal ciliary localization, consistent with the findings reported in *R26Arl13b-Fucci2a^Tg/Tg^* MEFs (Ford et al., 2018).

**Figure 2.**
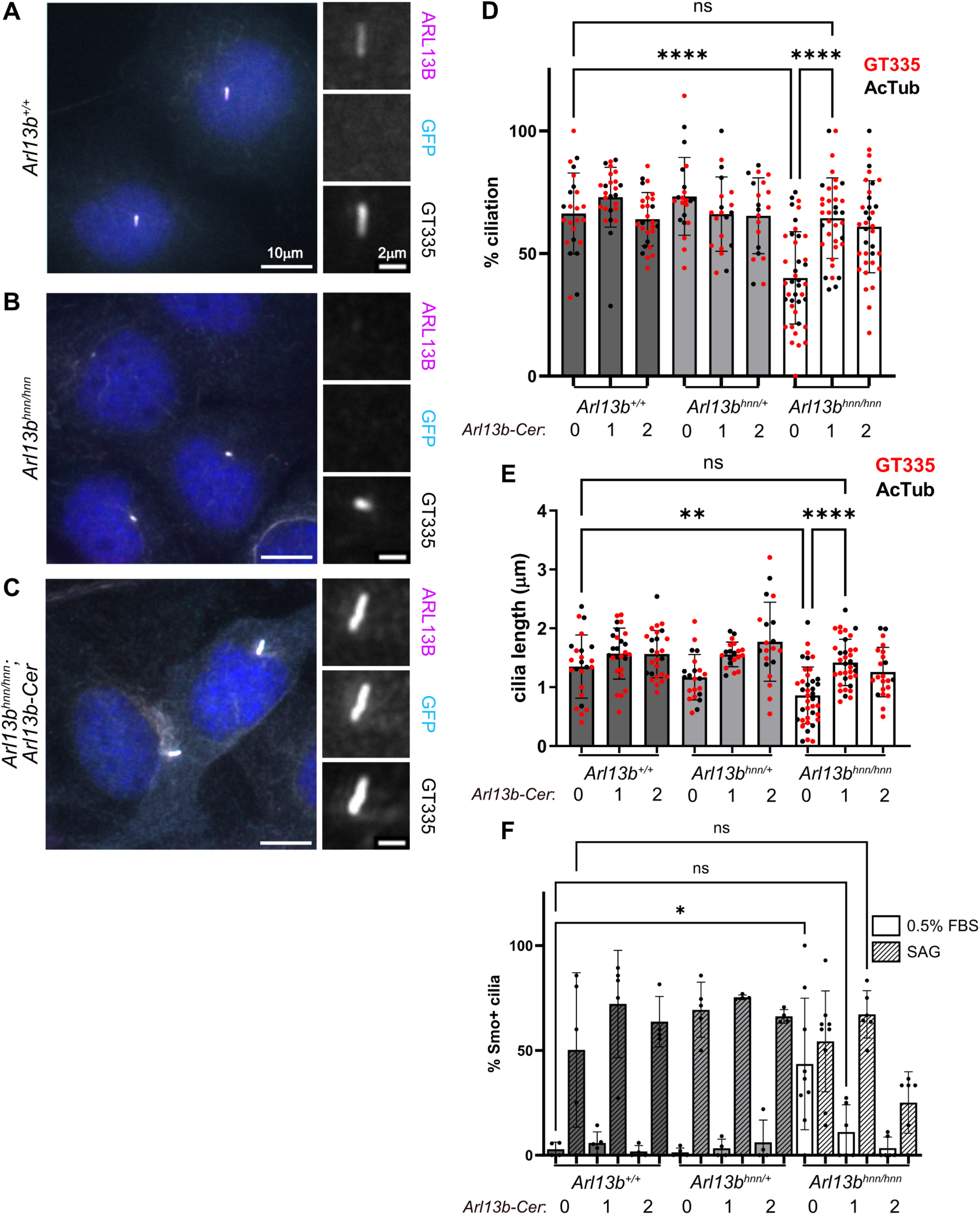
*Arl13b-Cerulean* rescues *Arl13b^hnn/hnn^* MEF ciliary phenotypes. Immortalized and serum-starved **A)** *Arl13b^+/+^*, **B)** *Arl13b^hnn/hnn^*, and **C)** *Arl13b^hnn/hnn^;Arl13b-Cerulean* MEFs stained with antibodies against ARL13B, GFP (recognizes Cerulean), and glutamylated tubulin GT335 (cilia marker). **D)** Percentage ciliated *Arl13b^+/+^*, *Arl13b^hnn/+^*, and *Arl13b^hnn/hnn^* MEFs without (0), or with (1, 2 copies) *Arl13b-Cerulean*, as indicated. One-way ANOVA with Tukey’s multiple comparisons test; adjusted *p* values: ns, not significant, *p* > 0.99; \*\*\*\**p* < 0.0001. None of the ciliation rates were significantly different when comparing cells with one or two copies of *Arl13b-Cerulean*, *p* > 0.52. **E)** Ciliary length based on the tubulin channel (glutamylated, red or acetylated, black). One-way ANOVA with Tukey’s multiple comparisons test; adjusted *p* values: ns, not significant, *p* > 0.92; \*\**p* < 0.01; \*\*\*\**p* < 0.0001. None of the cilia length measurements were significantly different when comparing cells with one or two copies of *Arl13b-Cerulean*, *p* > 0.85. **F)** Percent SMO-positive cilia observed under unstimulated (0.5% FBS, open bars) or stimulated (SAG, patterned bars) conditions. Two-way ANOVA with Tukey’s multiple comparisons test; adjusted *p* values: ns, not significant, *p* > 0.98; \**p* < 0.05. None of the SMO-positive percentages were significantly different when comparing cells with one or two copies of *Arl13b-Cerulean, p* > 0.99, except for the *Arl13b^hnn/hnn^* MEFs, *p* = 0.014. In the graphs, each point represents data from a single field of view with at least 150 total cilia examined for each genotype. Each MEF line was derived from a single embryo, and experiments were repeated three times.

To test whether ARL13B-Cerulean rescued the ciliogenesis defects in *Arl13b^hnn/hnn^*, we quantified the ciliation rate and cilia length in the MEF lines using antibodies against glutamylated and acetylated tubulin. We saw that the percentage of ARL13B-positive cilia was restored to wild-type levels in *Arl13b^hnn/hnn^;Arl13b-Cerulean* cells and that the zygosity of *Arl13b-Cerulean* did not significantly change the percentage of GFP-positive cilia (Fig. S2A,B). We observed a 25% decrease in the frequency of ciliated cells (from 66.0 to 40.7%) and a 33.5% decrease in average cilia length (from 1.40 to 0.93 microns) in the *Arl13b^hnn/hnn^* MEFs compared to wild type (Fig. 2D,E). In contrast, we found 63.1% of *Arl13b^hnn/hnn^;Arl13b-Cerulean* cells were ciliated, with a mean cilia length of 1.39 microns, similar to wild-type MEFs (Fig. 2D,E). When quantified separately, we saw no significant differences in ciliary length between glutamylated tubulin and acetylated tubulin antibodies (Fig. S2C). These data indicate that ARL13B-Cerulean rescues ciliation frequency and cilia length in *Arl13b^hnn/hnn^* MEFs.

The viability of *Arl13b^hnn/hnn^;Arl13b-Cerulean* mice suggested that Hh signaling was normal in these animals. To investigate this directly, we examined *Arl13b^hnn/hnn^;Arl13b-Cerulean* MEFs to ascertain whether ARL13B-Cerulean rescued the abnormal ciliary localization of Smoothened (SMO), the obligate transducer of the Hedgehog signaling pathway. We determined the percentage of SMO-positive cilia in the MEF lines under control and stimulated conditions. As expected, in unstimulated *Arl13b^hnn/hnn^* MEFs, we observed a 40% increase in the frequency of SMO-positive cilia compared to wild-type MEFs (Fig. 2F) (Larkins et al., 2011). In contrast, we found *Arl13b^hnn/hnn^;Arl13b-Cerulean* MEFs did not exhibit SMO-positive cilia, similar to wild-type MEFs cells (Fig. 2F). Furthermore, when we stimulated the cells for 24 hr with SMO agonist (SAG), we saw that all cell lines displayed an increased percentage of SMO-positive cilia (Fig. 2F). These data indicate that ARL13B-Cerulean restores normal SMO ciliary trafficking, consistent with a rescue of Hh signaling.

To determine the relative ARL13B and ARL13B-Cerulean protein expression levels in the immortalized MEFs, we performed western blots on whole cell lysates. We detected ARL13B-Cerulean only in cell lines that carried the biosensor (Fig. S2D). In contrast to our findings in embryo lysates, we observed similar levels of ARL13B-Cerulean and ARL13B expression in the MEFs (Fig. S2E). Again, we observed bands of the endogenous ARL13B size in *Arl13b^hnn/hnn^;Arl13b-Cerulean* cell lines, consistent with cleavage of the fusion protein (Fig. S2D).

To examine the relative transcript levels of *Arl13b* and *Arl13b-Cerulean*, we performed qRT-PCR using primers that differentiate between the 3’ ends of the transcripts (Fig. S2F). We detected *Arl13b-Cerulean* only in samples carrying the biosensor (Fig. S2G). In addition, we observed more *Arl13b-Cerulean* transcript in *Arl13b^+/+^* or *Arl13b^hnn/hnn^* samples with two copies of the biosensor compared to those with one copy (Fig. S2G). Despite these trends, increased amounts of mRNA or protein do not correlate with changes in ciliation rate or cilia length in the immortalized MEFs (Fig. 2D,E). Taken together, these data indicate that *Arl13b-Cerulean* rescues the *Arl13b^hnn/hnn^* cilia phenotypes, and the zygosity of the biosensor does not impact the rescue.

### Arl13b^hnn/hnn^;Arl13b-Cerulean kidneys do not develop cysts

As ciliary ARL13B regulates kidney development, we examined *Arl13b^hnn/hnn^;Arl13b-Cerulean* kidneys. We saw no change in the gross morphology of adult *Arl13b^hnn/hnn^;Arl13b-Cerulean* kidneys of either sex compared to controls (Fig. 3A). When we measured total kidney weight to body weight ratios, we observed a slight increase in male *Arl13b^hnn/hnn^;Arl13b-Cerulean* mice heterozygous for the biosensor (Fig. 3B, S3A). There was no difference in kidney weight to body weight ratio in females (Fig. 3C, S3B). We also performed histology to assess tissue structure and immunofluorescence staining to monitor cilia. We saw normal kidney architecture and ARL13B-positive cilia in *Arl13b^+/+^* and *Arl13b^hnn^*^/^*^+^;Arl13b-Cerulean* mice (Fig. 3D,E). In contrast to *Arl13b^V358A/V358A^* mice that express cilia-excluded ARL13B^V358A^ and develop kidney cysts by four weeks of age, adult *Arl13b^hnn/hnn^;Arl13b-Cerulean* mice did not have cystic kidneys (Fig. 3F) (Van Sciver et al., 2023). Of note, we found increased intra- and inter-tubule dilations in *Arl13b^hnn/hnn^;Arl13b-Cerulean* kidneys, which may be due to fibrosis, interstitial edema, or inflammation (Fig. 3F). To investigate whether fibrosis was contributing to this phenotype, we performed Sirius Red staining on kidney sections. We found no increased collagen staining in *Arl13b^hnn/hnn^;Arl13b-Cerulean* kidneys, indicating fibrosis did not underlie the dilations (Fig. S3C-E). We observed ARL13B-positive cilia in kidney tubules of all three genotypes (Fig. 3D-F). Altogether, these results indicate ARL13B-Cerulean is sufficient to support normal kidney ciliation and morphology.

**Figure 3.**
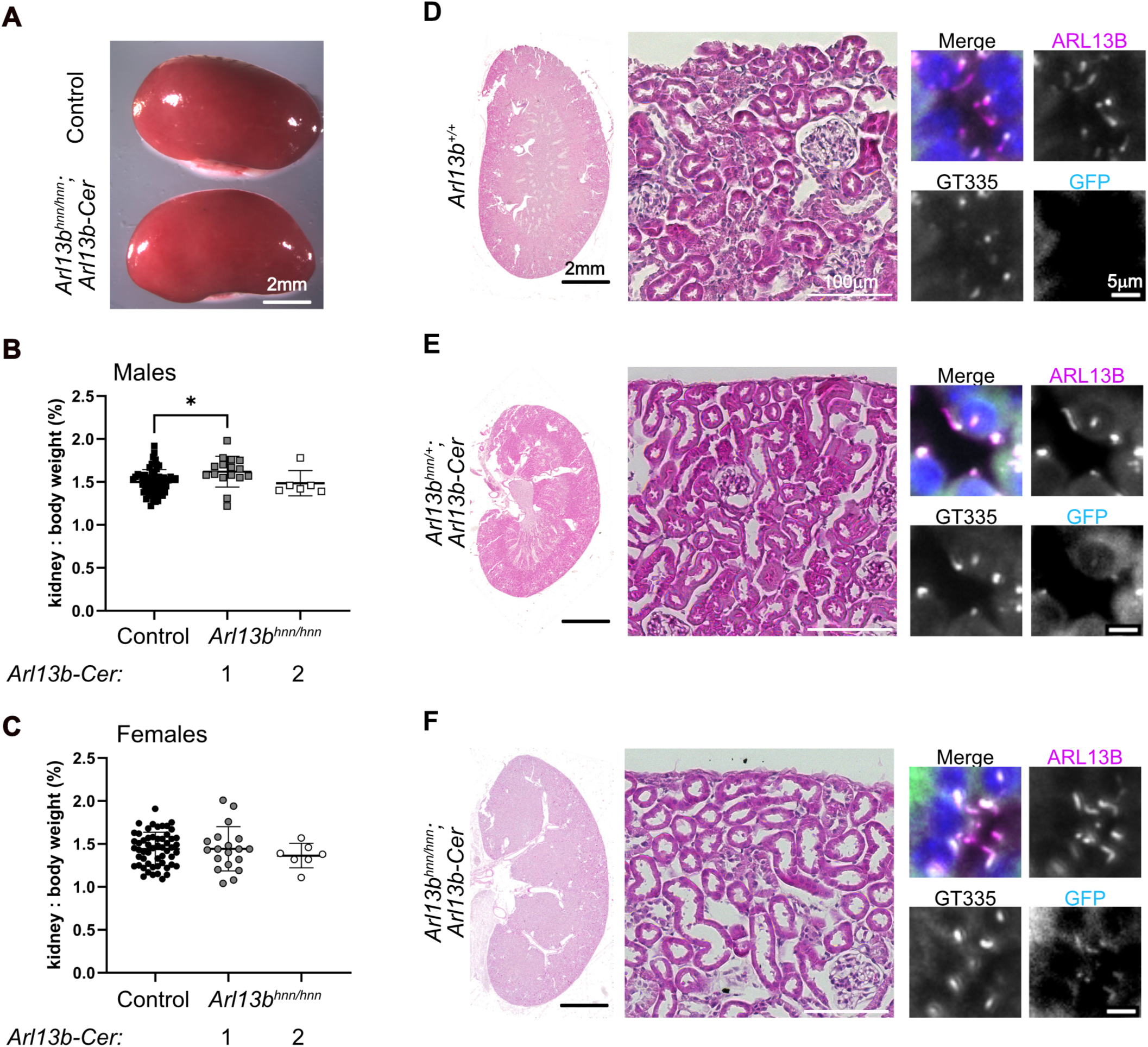
*Arl13b^hnn/hnn^;Arl13b-Cerulean* kidneys do not develop cysts. **A)** Gross morphology of adult control and rescue kidneys. **B)** Kidney weight as a percentage of body weight for males: control (black, *n* = 64), *Arl13b^hnn/hnn^;Arl13b-Cerulean* with one copy of *Arl13b-Cerulean* (gray, *n* = 16), *Arl13b^hnn/hnn^;Arl13b-Cerulean* with two copies of *Arl13b-Cerulean* (white, *n* = 6). Welch’s *t*-test showed a significant increase in the kidney weight of male mice carrying one copy of the biosensor compared to controls, \**p* < 0.05. **C)** Kidney weight as a percentage of body weight for females: control (black, *n* = 55), *Arl13b^hnn/hnn^;Arl13b-Cerulean* with one copy of *Arl13b-Cerulean* (gray, *n* = 19), *Arl13b^hnn/hnn^;Arl13b-Cerulean* with two copies of *Arl13b-Cerulean* (white, *n* = 7). Welch’s *t*-test showed no significant change in kidney weight of female mice carrying *Arl13b-Cerulean* compared to controls, *p* > 0.23. **D)** *Arl13b^+/+^*, **E)** *Arl13b^hnn/+^;Arl13b-Cerulean*, and **F)** *Arl13b^hnn/hnn^;Arl13b-Cerulean* kidney sections from adult mice stained with hematoxylin-eosin or antibodies against ARL13B, GFP, and glutamylated tubulin GT335. These images are representative of observed kidneys from at least three animals of each genotype.

### Cerebellar patterning is normal in *Arl13b^hnn/hnn^;Arl13b-Cerulean* mice

To determine whether ARL13B-Cerulean is functional in the brain, we examined *Arl13b^hnn/hnn^;Arl13b-Cerulean* brain morphology, focusing on the cerebellum. We performed hematoxylin-eosin staining on cerebellar sections, and found patterning and size were comparable between control (*Arl13b^hnn/+^*) and *Arl13b^hnn/hnn^;Arl13b-Cerulean* mice (Fig. 4A). ARL13B-positive cilia were present at the Purkinje cell layer (PCL) interface between the outer molecular layer (ML) and the inner granule layer (IGL) in *Arl13b^+/+^*, *Arl13b^hnn/+^;Arl13b-Cerulean* and *Arl13b^hnn/hnn^;Arl13b-Cerulean* cerebella (Fig. 4B). To determine the ARL13B and ARL13B-Cerulean protein levels in cerebella, we performed immunoblot analysis on lysates from adult tissues (Fig. S4A). We detected more ARL13B-Cerulean protein than endogenous ARL13B protein in cerebella that carried the biosensor (Fig. S4B). In addition, we detected increased amounts of ARL13B-Cerulean protein correlated with the zygosity of *Arl13b-Cerulean* (Fig. S4C). We saw no overt hydrocephaly in *Arl13b^hnn/hnn^;Arl13b-Cerulean* mice, although we noted slight enlargement of the lateral ventricle lumens compared to control animals (Fig. S4D,E). Choroid plexus cilia in *Arl13b^hnn/hnn^;Arl13b-Cerulean* brains were visible with antibodies against ARL13B or GFP (Fig. 4C). These findings demonstrate that brain development is grossly normal in *Arl13b^hnn/hnn^;Arl13b-Cerulean* animals.

**Figure 4.**
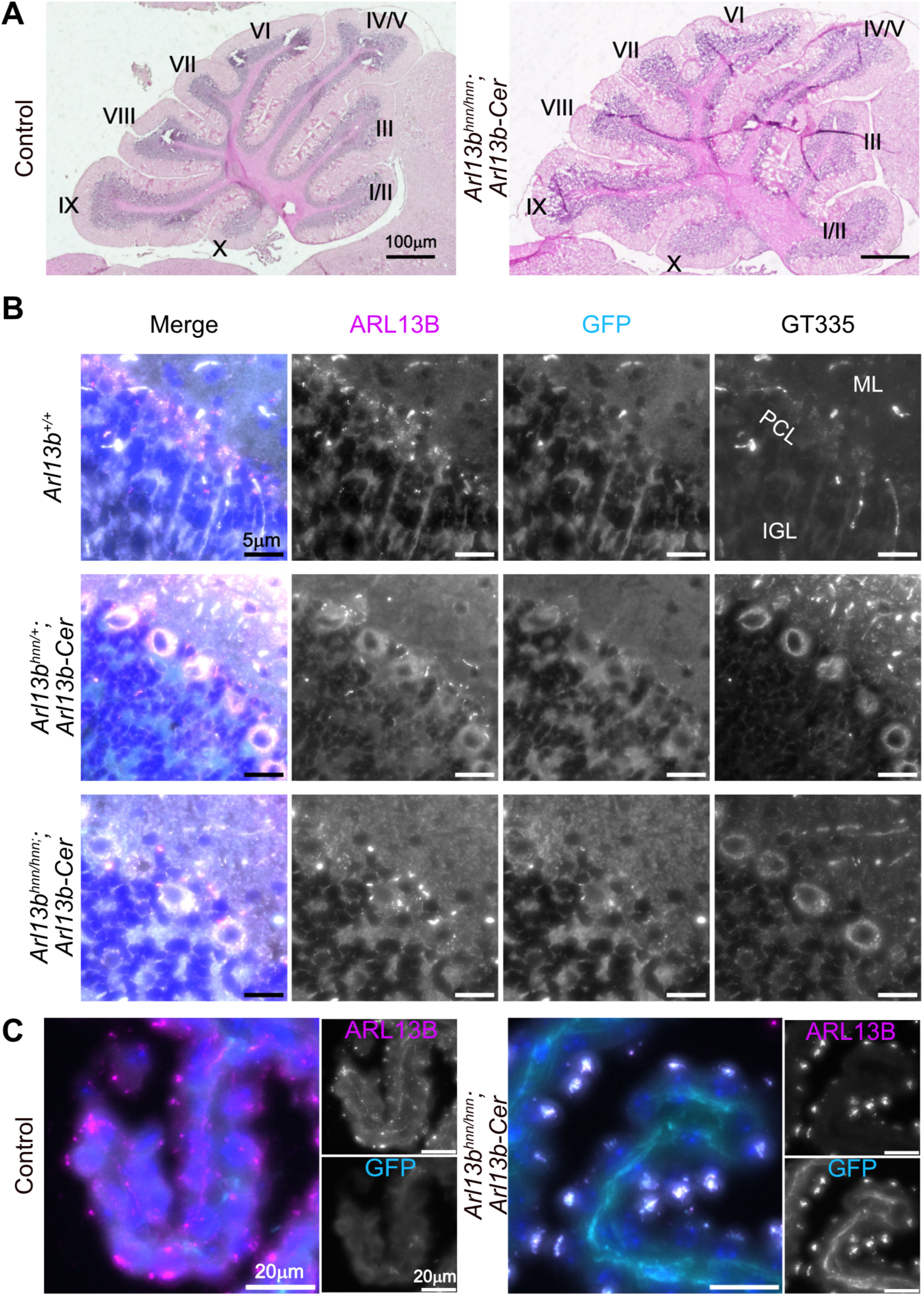
Cerebellar patterning is normal in *Arl13b^hnn/hnn^;Arl13b-Cerulean* mice. **A)** Sagittal sections of adult control and *Arl13b^hnn/hnn^;Arl13b-Cerulean* cerebella stained with hematoxylin-eosin. Roman numerals indicate major folia. **B)** ARL13B, GFP, and glutamylated tubulin GT335 staining of cilia in the Purkinje cell layer (PCL) of the cerebellum. ML (molecular layer), IGL (inner granule layer). **C)** Choroid plexus of control and *Arl13b^hnn/hnn^;Arl13b-Cerulean* brains stained with ARL13B and GFP antibodies.

### Pancreatic islet cilia are easily identified in *Arl13b^hnn/hnn^;Arl13b-Cerulean* mice

We examined the pancreas, a ciliated organ where ARL13B is expressed (Cho et al., 2022; Li et al., 2021; Li et al., 2022). We evaluated *Arl13b^hnn/hnn^;Arl13b-Cerulean* tissue morphology in sections stained with hematoxylin-eosin. We saw an increase in interstitial space surrounding the acinar cells of the *Arl13b^hnn/hnn^;Arl13b-Cerulean* pancreas compared to the control (Fig. 5A). We performed immunofluorescent staining and detected glucagon-producing alpha and insulin-producing beta cells normally arranged in islets of *Arl13b^hnn/hnn^;Arl13b-Cerulean* mice (Fig. 5B). We also observed ARL13B-positive cilia in islets from each of the groups, with overlapping GFP-positive staining in mice expressing ARL13B-Cerulean (Fig. 5C). These data indicate that at a gross level, the pancreas develops normally in *Arl13b^hnn/hnn^;Arl13b-Cerulean* mice.

**Figure 5.**
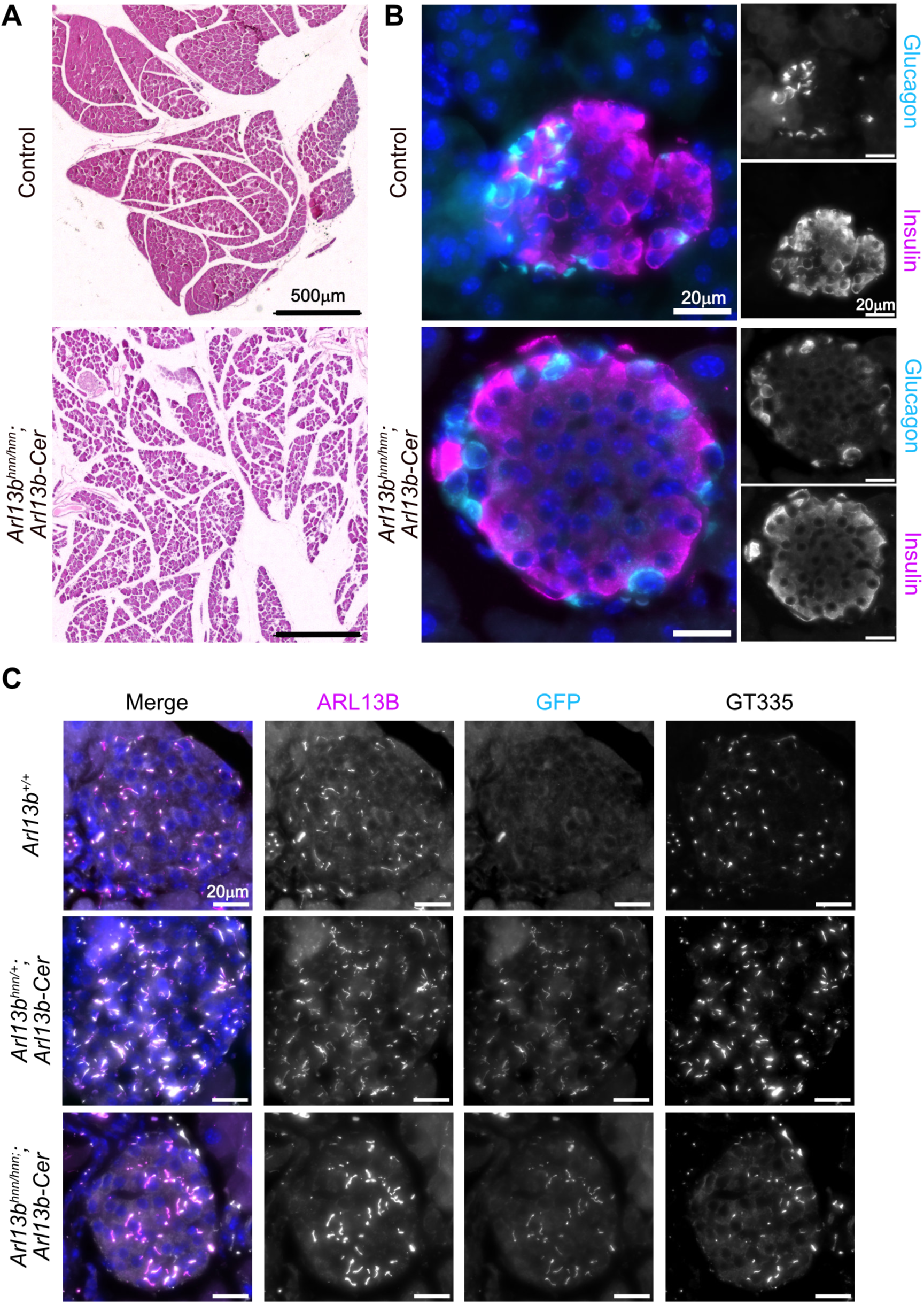
*Arl13b^hnn/hnn^;Arl13b-Cerulean* pancreatic islets appear normal. **A)** Pancreas sections from adult control or *Arl13b^hnn/hnn^;Arl13b-Cerulean* mice stained with hematoxylin-eosin. **B)** Immunofluorescent staining of islet cells in control and *Arl13b^hnn/hnn^;Arl13b-Cerulean* pancreas sections using antibodies against glucagon and insulin. **C)** Antibody staining of pancreatic islets showing ciliary ARL13B, GFP, and glutamylated tubulin GT335.

### *Arl13b^hnn/hnn^;Arl13b-Cerulean* males are infertile due to the absence of mature sperm

In the course of phenotyping, we mated *Arl13b^hnn/hnn^;Arl13b-Cerulean* with control *Arl13b^hnn/+^;Arl13b-Cerulean* mice. Females of either genotype and male control *Arl13b^hnn/+^;Arl13b-Cerulean* mice produced litters of normal size (Fig. 6A,B). In contrast, *Arl13b^hnn/hnn^;Arl13b-Cerulean* males did not produce any litters during 6-8 weeks of mating even though we detected copulation plugs, indicating normal mating behavior (Fig. 6B). We examined testis weight to body weight ratios and found no significant differences between *Arl13b^hnn/hnn^;Arl13b-Cerulean* and control mice (Fig. 6C, S5A).

**Figure 6.**
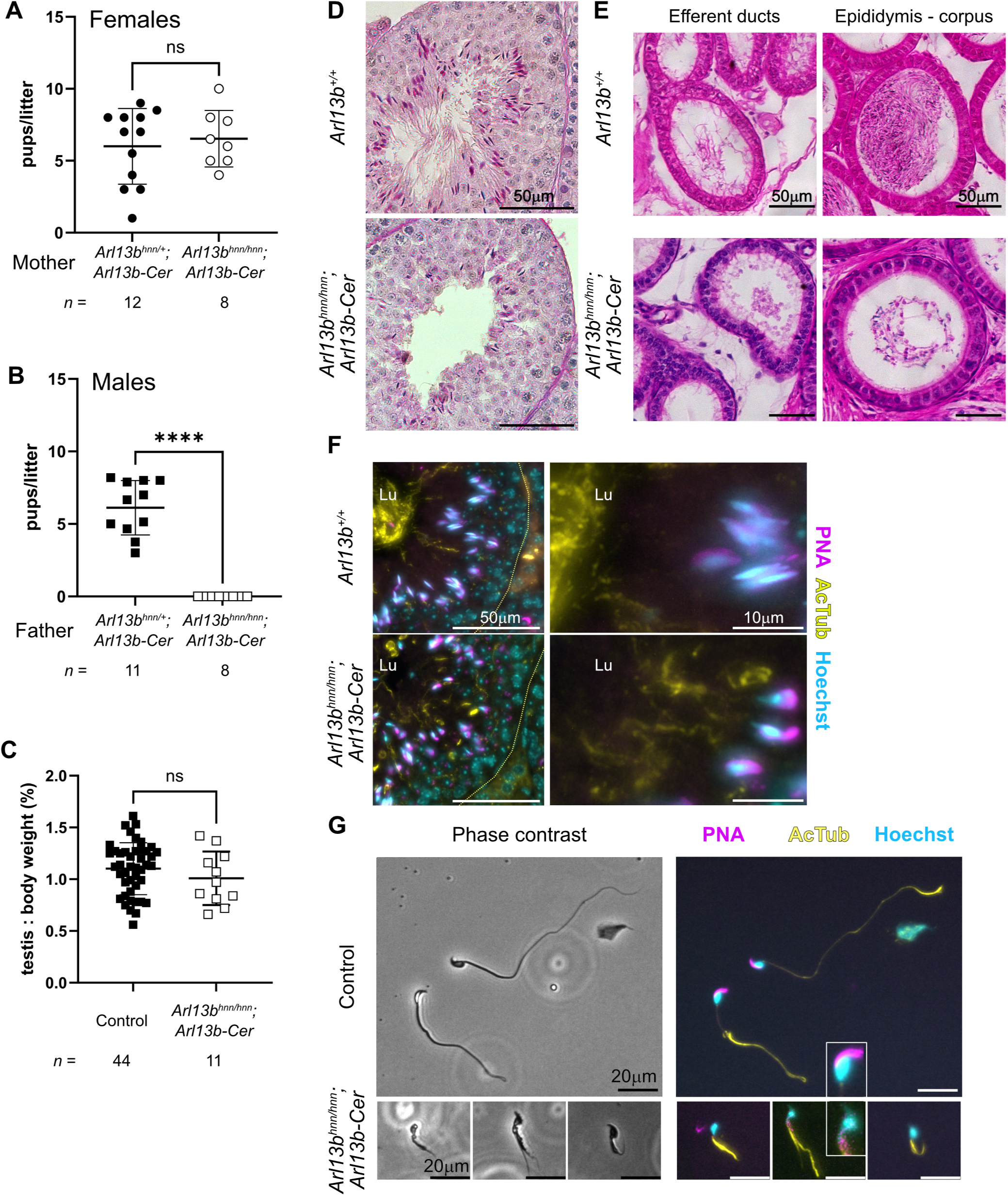
*Arl13b^hnn/hnn^;Arl13b-Cerulean* males are infertile. **A)** Average pups per litter when females of the indicated genotype were mated to control males (*Arl13b^hnn/+^;Arl13b-Cerulean*, *n* = 12 females tested: 181 pups from 29 litters; and *Arl13b^hnn/hnn^;Arl13b-Cerulean*, *n* = 6 heterozygous and 2 homozygous for *Arl13b-Cerulean* females tested: 122 pups from 18 litters). Mann Whitney test: ns, not significant, *p* > 0.95. **B)** Average pups per litter when males of the indicated genotype were mated to control females (*Arl13b^hnn/+^;Arl13b-Cerulean*, *n* = 11 males tested: 332 pups from 52 litters; and *Arl13b^hnn/hnn^;Arl13b-Cerulean*, *n* = 4 heterozygous and 4 homozygous for *Arl13b-Cerulean* males tested: 0 pups from 0 litters). Mann Whitney test: \*\*\*\**p*<0.0001. **C)** Testis weight as a percentage of body weight for control (*n* = 44) and *Arl13b^hnn/hnn^;Arl13b-Cerulean* (*n* = 8 heterozygous and 3 homozygous for *Arl13b-Cerulean*) male mice. Unpaired *t*-test: ns, not significant, *p* > 0.28. **D)** Testis sections from adult *Arl13b^+/+^* and *Arl13b^hnn/hnn^;Arl13b-Cerulean* males stained with PAS-H. **E)** Epididymis sections from adult *Arl13b^+/+^* and *Arl13b^hnn/hnn^;Arl13b-Cerulean* males stained with hematoxylin-eosin. **F)** Adult testis sections stained with peanut agglutinin lectin (PNA, acrosome) and antibodies against acetylated tubulin (AcTub, cilia) and Hoechst (nuclei). Dashed yellow lines indicate the basal lamina of the seminiferous tubule. Lu (lumen). Magnifications are shown on the right. **G)** Immunofluorescence of isolated control and *Arl13b^hnn/hnn^;Arl13b-Cerulean* sperm stained with PNA, AcTub, and Hoechst. Insets show magnified sperm heads.

To investigate the *Arl13b^hnn/hnn^;Arl13b-Cerulean* male infertility, we analyzed testis sections stained with periodic acid-Schiff reagent and hematoxylin (PAS-H). The overall *Arl13b^hnn/hnn^;Arl13b-Cerulean* seminiferous tubule architecture was normal (Fig. 6D).

The presence and patterning of germline (spermatogonia, spermatocytes) and support cells (Sertoli, Leydig) in *Arl13b^hnn/hnn^;Arl13b-Cerulean* testes did not differ from wild-type testes (Fig. 6D). We identified both round and elongating spermatids in the *Arl13b^hnn/hnn^;Arl13b-Cerulean* testis, consistent with normal meiosis (Fig. 6D). In later stages of spermatogenesis, elongating spermatids migrated toward the tubule lumen and sperm flagella were visible in the control testis (Fig. 6D). We had difficulty detecting mature sperm at the luminal surface of *Arl13b^hnn/hnn^;Arl13b-Cerulean* seminiferous tubules and did not observe sperm flagella in the lumen by histology (Fig. 6D). In addition, we were unable to visualize normal sperm by histology in the *Arl13b^hnn/hnn^;Arl13b-Cerulean* epididymis (Fig. 6E). Instead, the epididymides of *Arl13b^hnn/hnn^;Arl13b-Cerulean* mice appeared empty or contained debris (Fig. 6E). These data indicate the *Arl13b^hnn/hnn^;Arl13b-Cerulean* testis fails to produce mature flagellated sperm.

Nuclear condensation, acrosome formation, and flagellum biosynthesis occur during the final phase of spermatogenesis. We looked at nuclei (stained with Hoechst), the acrosome (marked with peanut agglutinin, PNA), and flagella (stained with antibodies against acetylated tubulin) in testis sections from wild-type and *Arl13b^hnn/hnn^;Arl13b-Cerulean* mice. We observed defects in the nucleus, acrosome, and flagellum morphology of *Arl13b^hnn/hnn^;Arl13b-Cerulean* sperm, suggesting defects in multiple aspects of sperm development (Fig. 6F). When we examined sections of the adult epididymis by immunofluorescence, we were able to identify the condensed nuclei of sperm heads and infrequent abnormal flagella in *Arl13b^hnn/hnn^;Arl13b-Cerulean* males (Fig. S5B). Taken together, these data indicate that some sperm, albeit abnormally formed sperm, are produced by the testes of the *Arl13b^hnn/hnn^;Arl13b-Cerulean* mice.

To examine spermatozoa head and tail morphology, we isolated sperm from wild-type, *Arl13b^hnn/+^;Arl13b-Cerulean* and *Arl13b^hnn/hnn^;Arl13b-Cerulean* epididymides. Overall, wild-type and control sperm displayed falciform heads with condensed nuclei and normal acrosomes as well as long flagella, whereas the few *Arl13b^hnn/hnn^;Arl13b-Cerulean* sperm we visualized exhibited abnormal heads and short, misshapen flagella (Fig. 6G). As we observed in the testis sections, the *Arl13b^hnn/hnn^;Arl13b-Cerulean* epididymal sperm displayed defects in acrosome formation and nuclear condensation (Fig. 6G). In addition, *Arl13b^hnn/hnn^;Arl13b-Cerulean* sperm tails were shorter than controls, with a mean length of 19.2 microns compared to 110.6 microns (Fig. 6G, S5C).

One explanation for the infertility and sperm morphology phenotype is that ARL13B-Cerulean expression interferes with normal ARL13B function despite not causing phenotypes in other tissues. However, we observed no phenotypic differences (reproductive or otherwise) in *Arl13b^+/+^* or *Arl13b^hnn/+^* males carrying one or two copies of *Arl13b-Cerulean* compared to those lacking the biosensor, arguing against such an effect (Fig. 6).

Conditional deletion of ARL13B in the efferent ducts, tubules connecting the testis to the epididymis, leads to defective cilia architecture, defective fluid resorption, and subfertility due to tubule occlusion (Augière et al., 2024). We investigated whether a similar phenotype was occurring in the *Arl13b^hnn/hnn^;Arl13b-Cerulean* males. Sections through the rete testis, where sperm accumulate before leaving the testis, showed normal morphology with no dilation of neighboring seminiferous tubules, suggesting that a blockage at this point is not the cause of infertility (Fig. S5D). When we directly examined the efferent ducts of *Arl13b^hnn/hnn^;Arl13b-Cerulean* males, we observed the presence of ARL13B-positive cilia on multiciliated epithelial cells (Fig. S5E). Therefore, ARL13B-Cerulean can replace the endogenous protein in the efferent ducts.

Taken together, our results show that ARL13B plays a critical role in sperm maturation and development. Furthermore, the male infertility phenotype suggests that ARL13B-Cerulean expression differs from that of endogenous ARL13B in a manner that renders ARL13B-Cerulean unable to fully compensate for endogenous ARL13B function in the male reproductive system.

### ARL13B-Cerulean is expressed in the testis and sperm

We investigated the expression levels of ARL13B and ARL13B-Cerulean in the testis via western blot. We observed that both proteins were present and that ARL13B is expressed at lower levels than ARL13B-Cerulean (Fig. 7A, S6A,B). To determine where ARL13B and ARL13B-Cerulean are expressed at the tissue level, we performed immunofluorescent staining on wild-type and *Arl13b^hnn/hnn^;Arl13b-Cerulean* testis sections. We detected a few ARL13B-positive cilia on cells located near the basal membrane of *Arl13b^+/+^* seminiferous tubules (Fig. 7B). In *Arl13b^hnn/hnn^;Arl13b-Cerulean* testis, we also identified these cilia with GFP antibody, indicating that both endogenous ARL13B and ARL13B-Cerulean were expressed in these support cells (Fig. 7B). The basal and myoid cells of *Arl13b^+/+^* and *Arl13b^hnn/hnn^;Arl13b-Cerulean* epididymides exhibited similar infrequent ciliary expression of ARL13B, consistent with published reports (Fig. S6C) (Bernet et al., 2018).

**Figure 7.**
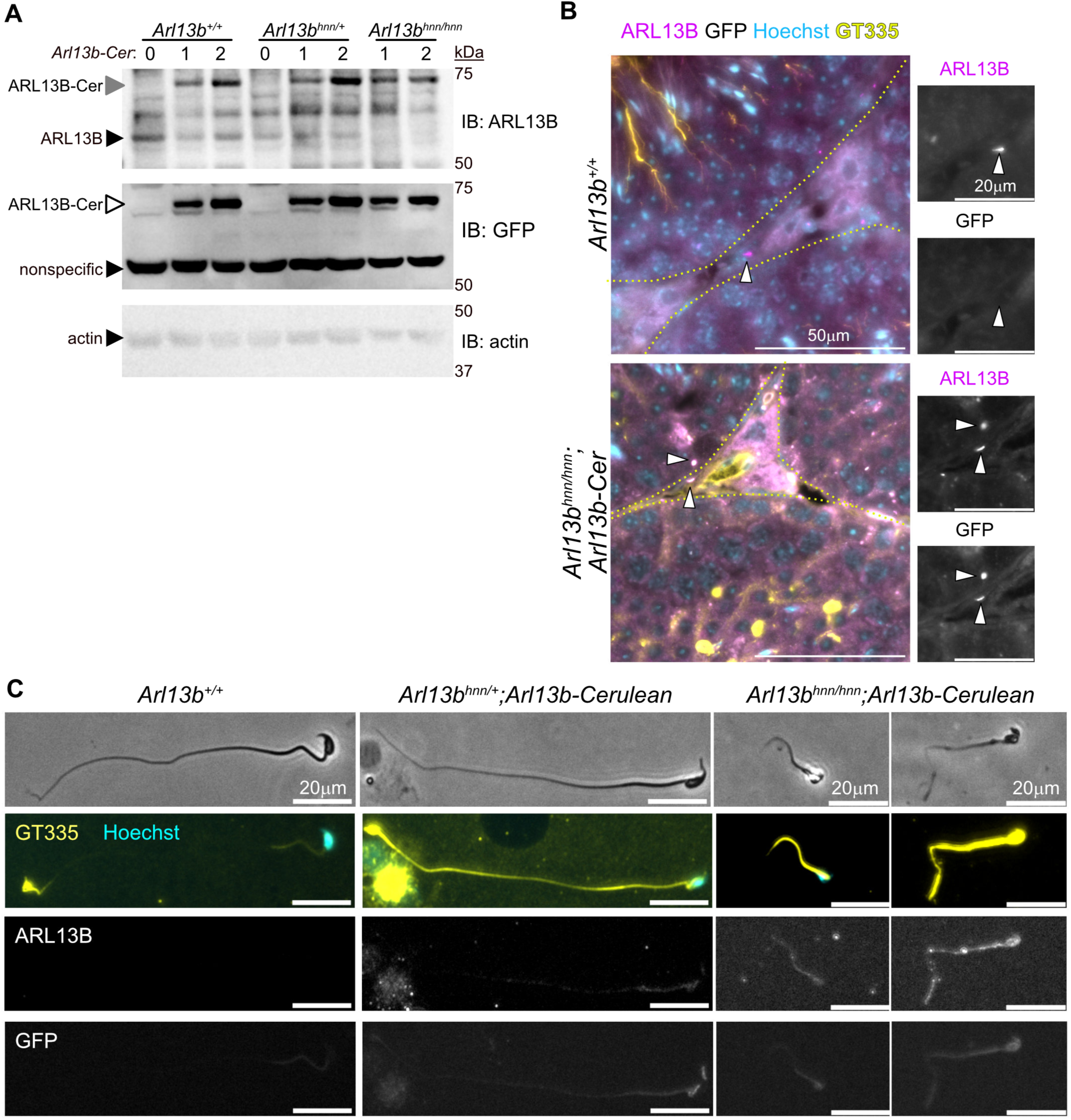
ARL13B-Cerulean is expressed in testis and sperm. **A)** Immunoblot (IB) of lysates from adult testes probed with antibodies against ARL13B, GFP, and actin. **B)** Testis sections from adult *Arl13b^+/+^* and *Arl13b^hnn/hnn^;Arl13b-Cerulean* males stained with antibodies against ARL13B, GFP, glutamylated tubulin GT335, and Hoechst. Dashed yellow lines indicate basal lamina of the seminiferous tubule. Magnifications are shown on the right. Arrowheads point to cilia. **C)** Sperm isolated from cauda epididymides of *Arl13b^+/+^*, *Arl13b^hnn/+^;Arl13b*-Cerulean, and *Arl13b^hnn/hnn^;Arl13b-Cerulean* males imaged by phase contrast and fluorescence microscopy using antibodies against glutamylated tubulin GT335, ARL13B, and GFP. Nuclei are stained with Hoechst.

To investigate the intracellular location of the ARL13B and ARL13B-Cerulean proteins, we stained isolated sperm with antibodies against ARL13B and GFP. We were unable to detect ARL13B in wild-type sperm by immunostaining, saw faint expression in the head and midpiece of *Arl13b^hnn/+^;Arl13b-Cerulean* sperm, and observed higher levels of expression along the whole length of *Arl13b^hnn/hnn^;Arl13b-Cerulean* sperm (Fig. 7C). When we stained with anti-GFP antibodies, we saw increased fluorescence of ARL13B-Cerulean from control *Arl13b^hnn/+^;Arl13b-Cerulean* to *Arl13b^hnn/hnn^;Arl13b-Cerulean* (Fig. 7C). Interestingly, we also saw increased tubulin glutamylation in sperm expressing *Arl13b-Cerulean* (Fig. 7C). Taken together, our data indicate that the ARL13B-Cerulean biosensor is functional in most tissues during development and reveals an essential role for ARL13B in spermiogenesis.

## DISCUSSION

Here, we showed that the ARL13B-Cerulean protein expressed by the *R26Arl13b-Fucci2aR* biosensor compensates for the loss of endogenous ARL13B in *Arl13b^hnn/hnn^* mice. We found that *Arl13b^hnn/hnn^;Arl13b-Cerulean* mice developed normally and survived into adulthood without gross morphological or behavioral defects indicating *R26Arl13b-Fucci2aR* is an inducible, functional *Arl13b* allele. Rescuing the embryonic lethality of *Arl13b^hnn/hnn^* allowed us to examine the function of ARL13B-Cerulean in several organs. We observed overall normal tissue anatomy and ciliation in the *Arl13b^hnn/hnn^;Arl13b-Cerulean* kidney, brain, and pancreas, although we did see a slight increase in interstitial spaces within these tissues. In addition, we examined the localization and expression level of ARL13B-Cerulean in these organs and detected normal ciliary enrichment and robust protein expression. Surprisingly, *Arl13b^hnn/hnn^;Arl13b-Cerulean* males were infertile. They exhibited low sperm counts and abnormal sperm shape, with short flagella and atypical acrosome structures. These findings suggest that ARL13B-Cerulean can functionally replace ARL13B in most developmental processes and reveal that endogenous ARL13B is required for normal spermatogenesis.

One unanticipated result of this work was that expression of ARL13B-Cerulean did not result in overt cilia phenotypes. We observed higher levels of ARL13B-Cerulean than ARL13B protein in tissue lysates but did not see significant length increases in MEF cilia or sperm flagella. This contrasts with previous findings *in vivo* where the *Arl13b* gene dosage directly correlates with ciliary length. Cilia are longer upon *Arl13b* overexpression in zebrafish or *Arl13b-mCherry* transgene expression in mice (Bangs et al., 2015; Lu et al., 2015; Pintado et al., 2017). Similarly, wild-type primary MEFs carrying two copies of the *Arl13b-Cerulean* biosensor display longer cilia (Ford et al., 2018). *In vivo*, wild-type mice homozygous for *Arl13b-Cerulean* have longer cilia in several tissues, although other tissues did not show this change (Ford et al., 2018). Another possibility is that cilia lengthening results from high levels of ARL13B. Perhaps the immortalization of MEFs in our study selected for ARL13B protein levels that do not reach this threshold. Furthermore, *in vivo Arl13b* overexpression has proven challenging: the broadly used *Arl13b-mCherry* transgenic mouse is the only line (of four founders) that has consistent expression, and the *Arl13b-EGFP* transgenic mouse is one of only two founders (Bangs et al., 2015; Delling et al., 2013). As transgenesis typically produces many more founders, the few ARL13B-expressing founders may reflect the fact that high ARL13B levels are not compatible with life. Thus, the protein levels of ARL13B-Cerulean may undergo similar selective pressure. In any case, ARL13B-Cerulean can functionally replace endogenous ARL13B with no adverse effects on organism-level growth and development.

This work reveals a novel function for ARL13B in spermatogenesis. ARL13B localizes to primary cilia of basal cells in the adult mouse epididymis and motile cilia in the efferent ducts to affect fluid flow and immune response in the male reproductive system (Augière et al., 2024; Bernet et al., 2018; Girardet et al., 2020; Girardet et al., 2022). Conditional deletion of ARL13B in the efferent ducts resulted in partial male infertility with defective cilia and improper tissue homeostasis (Augière et al., 2024). However, ARL13B is not known to function in the germline. We predicted that the spermatogenesis defect is due to abnormal ARL13B-Cerulean expression in the testis. We found that ARL13B-Cerulean is robustly expressed in the *Arl13b^hnn/hnn^;Arl13b-Cerulean* testis as well as in isolated sperm. Mice lacking ARL13B specifically in cilia, *Arl13b^V358A/V358A^*, are viable and fertile, so one possibility is that cellular ARL13B plays a critical function in spermatogenesis (Gigante et al., 2020). Answering this question will require using the *Arl13b^flox^* conditional null allele and appropriate Cre lines, which will also address issues of cell autonomy.

The cause of the defective spermatogenesis we observed in *Arl13b^hnn/hnn^;Arl13b-Cerulean* is unclear. It is unlikely that the biosensor had a neomorphic phenotype, as *Arl13b^+/+^* and *Arl13b^hnn/+^* males expressing *Arl13b-Cerulean* were fertile. Notably, the full-length *Arl13b* cDNA fused to Cerulean differs from endogenous ARL13B mRNA. Several proteins express distinct isoforms in somatic and germ cells (Eisa et al., 2021; Grassi et al., 2022; Konno et al., 2015; Nielsen and Raff, 2002). Perhaps specific ARL13B isoforms are expressed in the germline. Alternatively, the role of ARL13B in building a flagellum during spermatogenesis could be specific to flagellogenesis. While flagella and cilia possess axonemes, they also contain different components and perform distinct functions. For instance, mammalian sperm flagella have unique accessory structures such as the mitochondrial sheath, outer dense fibers, and the fibrous sheath. Our finding that the distribution of acetylation and glutamylation is different in sperm despite displaying relatively equivalent levels along the axonemes of MEF primary cilia is consistent with ARL13B’s role in these processes being tissue-dependent. Furthermore, ARL13B may normally interact with specific partners to contribute to the building of the flagellum, and ARL13B-Cerulean may be unable to replace endogenous protein for those interactions. Clearly, some difference between endogenous ARL13B and ARL13B-Cerulean is functionally relevant to spermatogenesis.

A key feature of the regulatory GTPase family to which ARL13B belongs is the ability to work with distinct effector proteins depending on cell type, subcellular location, or timing. ARL13B may have effectors specific to the testis or germline that play a role during spermatogenesis (nuclear condensation, acrosome formation, flagella outgrowth). On the other hand, some of ARL13B’s known effectors regulate spermatogenesis, suggesting the defects reflect a more general function of ARL13B. For example, ARL13B associates with the exocyst complex to regulate intracellular vesicle trafficking, with the BBSome to facilitate cargo coupling on the membrane, and with members of the IFT-B complex to regulate ciliary traffic (Barral et al., 2012; Cevik et al., 2013; Liu et al., 2023; Nozaki et al., 2017). Conditional deletion of *Exoc1*, encoding an exocyst component, in spermatogonia leads to spermatocyte aggregation and impaired fertility (Osawa et al., 2021). Mice carrying mutations in BBSome subunits exhibit male infertility due to an absence of sperm flagella (Davis et al., 2007; Mykytyn et al., 2004; Nishimura et al., 2004). Similarly, mutations in IFT proteins result in defective sperm morphology, quantity, and/or motility (Liu et al., 2017; San Agustin et al., 2015; Shi et al., 2019; Zhang et al., 2016; Zhang et al., 2017). Further work is needed to determine whether these effectors contribute to the mechanism(s) underlying the infertility we observed in the *Arl13b^hnn/hnn^;Arl13b-Cerulean* males.

In summary, our data show that full-length *Arl13b* cDNA with a Cerulean tag can functionally replace the endogenous *Arl13b* gene in several tissues and biological processes, with the notable exception of spermatogenesis. Future studies will determine the cell types that require ARL13B for spermatogenesis, as well as the mechanism by which ARL13B regulates spermatogenesis. Our findings expand upon previous work illustrating that the *R26Arl13b-Fucci2aR* biosensor is a powerful tool to monitor cilia and cell cycle progression. Our work is significant in demonstrating that *Arl13b-Cerulean* is a functional, inducible allele of *Arl13b*.

## MATERIALS AND METHODS

### Mice

All animal experiments were approved by the Institutional Animal Care and Use Committee of Emory University and were performed in compliance with guidelines established by the National Institute of Health. Mouse alleles used in this study were *Arl13b^hnn^* (MGI:3578151), *CMV-cre* (Tg(CMV-cre)1Cgn, MGI:2176180) acquired from the Jackson Laboratory (RRID:IMSR_JAX:006054), and *R26Arl13b-Fucci2aR* (Gt(ROSA)26Sor^tm1(CAG-Cerulean/Arl13b,-Venus/GMNN,-Cherry/CDT1)Rmort^, MGI:6193734) purchased from EMMA (EM:12168). Adult animals were over 8 weeks of age. Genotyping was performed on biopsied samples (ear punch or yolk sac) by Transnetyx using real-time PCR.

### Antibodies and reagents

Antibodies used in this study are listed in Table S1.

### Cell culture

MEFs were derived from E12.5 mouse embryos and cultured in DMEM supplemented with 10% fetal bovine serum (FBS) and 1% penicillin/streptomycin at 5% CO_2_ at 37°C. MEFs were immortalized by transfection with a plasmid containing the large T antigen of SV40. Cell lines were derived from a single embryo of each genotype. Data presented based on MEF experiments are representative (IF images) or cumulative (graphs) of three technical replicates.

For immunofluorescence, immortalized MEFs were plated on glass coverslips and serum-starved for 24 hours in DMEM supplemented with 0.5% FBS. Cells were fixed in 4% paraformaldehyde (PFA) for 10 minutes, washed three times with phosphate-buffered saline (PBS) for 5 minutes each, and blocked with tris buffered saline (TBS) + 5% goat serum + 0.1% Triton X-100 (blocking buffer) for 10 minutes. Cells were incubated with primary antibodies diluted in blocking buffer overnight at 4°C, washed three times with blocking buffer for 5 minutes each, and incubated with secondary antibodies diluted in blocking buffer for 1 hour at room temperature in the dark. Finally, cells were washed three times with blocking buffer for 5 minutes each and mounted onto glass slides using Prolong Gold.

To assess Hh response, immortalized MEFs were plated on glass coverslips and serum-starved for 24 hours in DMEM with 0.5% FBS. An additional 24 hours of treatment was performed using either 0.5% FBS DMEM alone or supplemented with 200nM SAG (Sigma Aldrich). Cells were fixed and stained as detailed above using antibodies against SMO (Ocbina et al., 2011) and cilia (glutamylated or acetylated tubulin).

Images were obtained with a BioTek Lionheart FX automated microscope (Agilent) and processed with Gen5 imaging software (version 3.11, Agilent) and FIJI (release 2.16.0) (Schindelin et al., 2012). For cilia counts, we quantified the number of nuclei and glutamylated or acetylated tubulin-positive cilia in each field of view with multiple fields imaged. To determine cilia length, the CiliaQ plugin for FIJI was used with measurements taken using the tubulin channel (Hansen et al., 2021).

### Western blot

Lysates were obtained by homogenizing tissue with Pierce RIPA Buffer (ThermoScientific) containing protease inhibitor cocktail (Sigma Aldrich) using a mechanical pestle. After sonication on ice, lysates were incubated at 4°C for 30 minutes turning end-over-end. Lysates were centrifuged at 13000 rpm for 15 minutes at 4°C and the protein content of the supernatant was determined using the Pierce BCA Protein Assay (ThermoScientific). Equal amounts (30µg) of protein were separated by SDS-PAGE and transferred onto nitrocellulose membranes. After blocking with StartingBlock T20 PBS (ThermoScientific) (antibody block) for at least 1 hour, membranes were incubated with primary antibodies diluted in antibody block for 1 hour at room temperature or overnight at 4°C. Primary antibodies were detected using fluorescently conjugated secondary antibodies. The Azure600 imaging system (Azure Biosystems) and FIJI software were used to visualize and quantify proteins, with anti-actin-rhodamine loading control used to normalize band intensity. Each series of lysates was run at least three times, and the presented images are representative of those repeats. Band intensity was measured using FIJI, and the graphed results indicate the average ± standard error.

### Histological analyses

Mice were perfused transcardially with 4% PFA in PBS. Organs were dissected, postfixed in 4% PFA (kidney, pancreas, brain) or Bouin’s fixative (testis) overnight at 4°C, then dehydrated in ethanol for paraffin embedding or cryoprotected with 30% sucrose in phosphate buffer and embedded in Tissue-Tek OCT compound (Sakura) for frozen sectioning. Paraffin sections (kidney, testis, pancreas) were acquired at 5 µm and frozen sections (brain) were acquired at 10 µm. Sections were mounted onto Superfrost Plus microscope slides.

For histology, deparaffinized or rehydrated frozen sections were stained with hematoxylin and eosin, Sirius Red and fast green, or periodic acid and Schiff’s reagent with hematoxylin (PAS-H) following standard protocols. Slides were coverslipped with Cytoseal 60 (Epredia). For immunofluorescence, antigen retrieval was performed on deparaffinized sections using hot 10 mM citrate buffer, pH 6.0, before incubation with primary antibodies overnight at 4°C. Sections were then incubated with fluorescent secondary antibodies and Hoechst, and coverslipped with Prolong Gold (ThermoFisher).

### Quantitative reverse transcriptase PCR (qRT-PCR)

Cells were collected, snap frozen, and stored at -80°C. RLT lysis buffer (Qiagen) was added, and RNA was purified with the RNeasy Mini Kit (Qiagen) according to the manufacturer’s protocol. Potential genomic DNA contamination was eliminated by incubation with RNase-free DNase (Qiagen). To generate cDNA, total RNA was reverse transcribed using iScript (Bio-Rad). The SsoAdvanced Universal SYBR Green Supermix kit (Bio-Rad) was used to perform real-time PCR on prepared cDNA. Each reaction was performed in triplicate for each biological sample and normalized to *Gapdh*. Primers were: *FucciQ “F3”*: 5’-GTGATGCTCAGGACACGATC-3’; *FucciQ “R3”*: 5’-CGGTGGTGCAGATGAACTTC-3’; *Arl13bQ “R4”*: 5’-TTGTCTTGCCCATCATCAGC-3’; *Gapdh-F*: 5’-CGTCCCGTAGACAAAATGGT-3’; *Gapdh-R*: 5’-GAATTTGCCGTGAGTGGAGT-3’.

### Sperm isolation and staining

The epididymis was dissected and kept in a Petri dish containing PBS. Sperm were released from the cauda epididymis by puncturing the tissue with fine forceps. The sperm suspension was placed on a glass-bottom 35 mm dish and incubated for 30 minutes in a CO_2_ incubator at 37°C. Next, the sperm cells were fixed in 4% PFA in PBS for 10 minutes at room temperature, permeabilized and blocked for 1 hour in TBS + 5% goat serum + 0.1% Triton X-100 (blocking buffer) and incubated with primary antibodies diluted in blocking buffer for 1 hour at room temperature. After washing three times with blocking buffer for 5 minutes each, cells were incubated with secondary antibodies diluted in blocking buffer for 1 hour at room temperature in the dark. Another three washes with blocking buffer for 5 minutes each were followed by the addition of glass coverslips using Prolong Gold as the mounting medium.

### Statistical analyses

Data were analyzed by the indicated statistical test using Prism version 10.5 (GraphPad). Summarized data are presented as the mean ± standard deviation, unless noted. Images are representative of a minimum of three biological samples and sample numbers for summarized data are indicated on the graphs and in the figure legends.

## Acknowledgments

We thank lab members for discussion and comments on the manuscript, Emory University’s NextGen 2024 summer high school students for piloting the histology staining procedures, and the Emory University Mouse Transgenic and Gene Targeting Core (RRID:SCR_023535) for technical assistance.

The cryopreserved B6;129P2-Gt(ROSA)26Sor^tm1(CAG-Cerulean/Arl13b,-Venus/GMNN,-Cherry/CDT1)Rmort^/H sperm was obtained from the Mary Lyon Centre at MRC Harwell which distributes this strain on behalf of the European Mouse Mutant Archive (EM:12168). These mice were originally produced at the University of Lancaster.

## Author contributions

Conceptualization: A.B.L., I.M.W., T.T.T., R.E.V.S., T.C.

Methodology: A.B.L., T.T.T., R.E.V.S.

Formal analysis: A.B.L., I.M.W., T.T.T.

Investigation: A.B.L., I.M.W., T.T.T.

Writing – original draft: A.B.L.

Writing – review & editing: A.B.L., I.M.W., T.T.T., R.E.V.S., T.C.

Visualization: A.B.L., T.T.T., R.E.V.S.

Supervision: T.C.

Project administration: T.C.

Funding acquisition: T.C.

## Data availability

All relevant data can be found within the article and its supplementary information.

---

**Table S1.** Antibodies used in this study.

| **Antibody** | **Manufacturer** | **Catalog number** | **Dilution**<br>**(purpose)** |
| --- | --- | --- | --- |
| Polyclonal rabbit anti-ARL13B | ProteinTech | 17711-1-AP | 1:2000 (IB)<br>1:1000 (IF) |
| Polyclonal chicken anti-GFP | Abcam | Ab13970 | 1:1000 (IF) |
| Mouse anti-glutamylated tubulin (GT335) | AdipoGen | AG-2032020 | 1:1000 (IF) |
| Monoclonal mouse anti-acetylated tubulin (AcTub) | Sigma | T6793 | 1:2500 (IF) |
| Polyclonal rabbit anti-SMO | Anderson lab | n/a | 1:1000 (IF) |
| Polyclonal rabbit anti-insulin | Cell Signaling | 3014 | 1:1000 (IF) |
| Monoclonal mouse anti-glucagon | Abcam | Ab10988 | 1:500 (IF) |
| PNA-rhodamine | Vector Labs | RL-1072 | 1:1000 (IF) |
| AlexaFluor-conjugated secondary | ThermoFisher | A11039, A31572, A21206, A11029, A21202 | 1:500 (IF) |
| Goat anti-mouse-AF647 | Jackson Immuno | 115-606-003 | 1:500 (IF) |
| Polyclonal rabbit anti-GFP | Abcam | Ab290 | 1:1000 (IB) |
| Actin-rhodamine | Bio-Rad | 12004163 | 1:5000 (IB) |
| IR680LT secondary | LI-COR | 926-68020 | 1:5000 (IB) |
| IR800CW secondary | LI-COR | 925-32213 | 1:5000 (IB) |

IF, immunofluorescence; IB, immunoblotting

**Figure S1. Weight analysis and quantification of embryonic protein expression.**

Body weights of control (black) and *Arl13b^hnn/hnn^;Arl13b-Cerulean* (white) **A)** male and **B)** female mice at the time of sacrifice. Nonlinear regression using the Gompertz growth model and least squares fit identified a single curve for each sex, indicating *Arl13b^hnn/hnn^;Arl13b-Cerulean* mice weights are not significantly different from controls: males *p* = 0.59, R^2^ = 0.82; females *p* = 0.14, R^2^ = 0.85. **C)** Quantification of signal intensity for embryo lysate western blots; endogenous ARL13B (black bars) and ARL13B-Cerulean (gray bars). Data are normalized to actin and set relative to the ARL13B band intensity in wild-type embryos. Bars represent the mean ± standard error of technical replicates. One-way ANOVA for ARL13B protein levels, *p* = 0.001, and for ARL13B-Cerulean protein levels, *p* = 0.019.

**Figure S2. Characterization of MEF lines.**

**A)** Percent ARL13B-positive and **B)** GFP-positive cilia for *Arl13b^+/+^*, *Arl13b^hnn/+^*, and *Arl13b^hnn/hnn^* cells without (0), or with (1, 2 copies) *Arl13b-Cerulean*, as indicated. One-way ANOVA with Tukey’s multiple comparisons test; adjusted *p* values: ns, not significant, *p* > 0.99, \*\*\**p* = 0.0001, \*\*\*\**p* < 0.0001. None of the ARL13B-positive rates were significantly different when comparing cells with one or two copies of *Arl13b-Cerulean*, *p* > 0.09. **C)** Comparison of ciliary length determined using antibodies against glutamylated tubulin GT335 (red circles) or acetylated tubulin (black circles) in *Arl13b^+/+^*, *Arl13b^hnn/+^*, and *Arl13b^hnn/hnn^* cells without (0), or with (1, 2 copies) *Arl13b-Cerulean*, as indicated. Two-way ANOVA with Tukey’s multiple comparisons test: overall interaction between genotype and antibody was not significant, *p* = 0.81; adjusted *p* values: ns, not significant, *p* > 0.05; \**p* < 0.05, \*\**p* < 0.01. None of the length measurements (using the same antibody, Actub or GT335), were significantly different when comparing cells with one or two copies of *Arl13b-Cerulean*, *p* > 0.96. Each point represents data from a single field of view with at least 150 cilia examined for each genotype and each experiment. **D)** Immunoblot analysis of MEF lysates probed with antibody against ARL13B (upper image) or actin (lower image). **E)** Quantification of signal intensities for ARL13B (black bars) and ARL13B-Cerulean (gray bars), with data normalized to actin and set relative to the ARL13B band intensity in wild-type MEFs. Bars represent the mean ± standard error of technical replicates. One-way ANOVA for ARL13B protein levels, *p* = 0.003, and for ARL13B-Cerulean protein levels, *p* = 0.0004. **F)** Schematic showing locations of primers used in qRT-PCR. **G)** Quantification of endogenous *Arl13b* (black) and *Arl13b-Cerulean* (gray) transcript levels in cell lines, normalized to *Gapdh* and set relative to *Arl13b* expression in wild-type MEFs. The graph shows the mean ± standard deviation for three technical replicates. One-way ANOVA for *Arl13b* transcript levels, *p* < 0.0001, and for *Arl13b-Cerulean* transcript levels, *p* <0.0001.

**Figure S3. *Arl13b^hnn/hnn^;Arl13b-Cerulean* kidneys are not fibrotic.**

Kidney weight versus body weight of control (black) and *Arl13b^hnn/hnn^;Arl13b-Cerulean* (white) **A)** male and **B)** female mice at time of sacrifice. Nonlinear regression using a straight line model and least squares fit identified a single curve for each sex, indicating the kidney:body weight ratios of rescue mice are not significantly different from controls: males *p* = 0.22, R^2^ = 0.92; females *p* = 0.40, R^2^ = 0.82. **C-E)** Sirius red-fast green staining of **C)** *Arl13b^+/+^*, **D)** *Arl13b^hnn/+^;Arl13b-Cerulean*, and **E)** *Arl13b^hnn/hnn^;Arl13b-Cerulean* kidney sections. Reddish-purple color indicates collagen associated with fibrosis, and green shows counterstain.

**Figure S4. Enlarged ventricles in *Arl13b^hnn/hnn^;Arl13b-Cerulean* mice.**

**A)** Immunoblot (IB) showing lysates from cerebella of adult mice probed with antibodies against ARL13B and GFP. Actin loading control is shown below. **B)** Quantification of signal intensities for the blots probed with ARL13B antibody, showing ARL13B (black bars) and ARL13B-Cerulean (gray bars) with data normalized to actin and relative to the ARL13B band intensity in wild-type cerebellum. Bars represent the mean ± standard error of technical replicates. One-way ANOVA for ARL13B protein levels, *p* = 0.09, and for ARL13B-Cerulean protein levels, *p* = 0.11. **C)** Quantification of ARL13B-Cerulean signal intensity for the blots probed with GFP antibody. Values are normalized to actin and relative to the ARL13B-Cerulean band intensity in wild-type cerebellum. Bars represent the mean ± standard error of technical replicates. One-way ANOVA for ARL13B-Cerulean protein levels, *p* = 0.03. **D)** Coronal sections through the lateral ventricles of adult control and *Arl13b^hnn/hnn^;Arl13b-Cerulean* brains stained with hematoxylin-eosin. **E)** Ventricular area as a percentage of total brain area. Each point represents a measured brain section; control *n* = 3, *Arl13b^hnn/hnn^;Arl13b-Cerulean n* = 5. Mann Whitney test, \**p* = 0.015.

**Figure S5. *Arl13b^hnn/hnn^;Arl13b-Cerulean* males have normal testis and epididymis morphology.**

**A)** Testis weight versus body weight of control (black) and *Arl13b^hnn/hnn^;Arl13b-Cerulean* (white) males. Nonlinear regression using a straight line model and least squares fit identified a single curve for each genotype, indicating the testis:body weight ratios of rescue males are not significantly different from controls: *p* = 0.99, R^2^ = 0.85. **B)** Immunofluorescence of control and *Arl13b^hnn/hnn^;Arl13b-Cerulean* corpus epididymis sections stained with glutamylated tubulin GT335 and Hoechst. **C)** Measurement of cauda epididymis-isolated sperm tail length, with at least 65 spermatozoa measured per genotype, including samples from at least three males per genotype. Unpaired *t*-test: \*\*\*\**p* < 0.0001. **D)** Section through the rete testis of *Arl13b^hnn/hnn^;Arl13b-Cerulean* adult stained with hematoxylin-eosin. **E)** Efferent duct sections stained with antibodies against glutamylated tubulin (GT335, white), acetylated tubulin (AcTub, white), GFP (white), ARL13B (magenta), and Hoechst (cyan). The final panel shows the ARL13B channel alone. Ductal lumens are to the right in each image.

**Figure S6. Expression of ARL13B-Cerulean and ARL13B in testis and epididymis.**

**A)** Quantification of signal intensity for ARL13B western blots of testis lysates with data normalized to actin and set relative to the ARL13B band intensity in wild-type testis. Bars represent the mean ± standard error of technical replicates. One-way ANOVA for ARL13B protein levels, *p* = 0.52, and for ARL13B-Cerulean protein levels, *p* = 0.013. **B)** Quantification of ARL13B-Cerulean signal intensity for the blots probed with GFP antibody. Values are normalized to actin and relative to the ARL13B-Cerulean band intensity in wild-type testis. Bars represent the mean ± standard error of technical replicates. One-way ANOVA for ARL13B-Cerulean protein levels, *p* = 0.046. **C)** Epididymis sections from adult *Arl13b^+/+^* and *Arl13b^hnn/hnn^;Arl13b-Cerulean* males stained with antibodies against ARL13B, GFP, glutamylated tubulin GT335, and Hoechst. Arrowheads point to cilia.

## REFERENCES

Augière, C., Campolina-Silva, G., Vijayakumaran, A., Medagedara, O., Lavoie-Ouellet, C., Joly Beauparlant, C., Droit, A., Barrachina, F., Ottino, K., Battistone, M. A., et al. (2024). ARL13B controls male reproductive tract physiology through primary and motile cilia. Commun. Biol. 7, 1318.

Bangs, F. K., Schrode, N., Hadjantonakis, A.-K. and Anderson, K. V. (2015). Lineage specificity of primary cilia in the mouse embryo. Nat. Cell Biol. 17, 113–122.

Barral, D. C., Garg, S., Casalou, C., Watts, G. F. M., Sandoval, J. L., Ramalho, J. S., Hsu, V. W. and Brenner, M. B. (2012). Arl13b regulates endocytic recycling traffic. Proc. Natl. Acad. Sci. 109, 21354–21359.

Bay, S. N., Long, A. B. and Caspary, T. (2018). Disruption of the ciliary GTPase Arl13b suppresses Sonic hedgehog overactivation and inhibits medulloblastoma formation. Proc. Natl. Acad. Sci. 115, 1570–1575.

Bernet, A., Bastien, A., Soulet, D., Jerczynski, O., Roy, C., Bianchi Rodrigues Alves, M., Lecours, C., Tremblay, M.-È., Bailey, J. L., Robert, C., et al. (2018). Cell-lineage specificity of primary cilia during postnatal epididymal development. Hum. Reprod. 33, 1829–1838.

Borovina, A., Superina, S., Voskas, D. and Ciruna, B. (2010). Vangl2 directs the posterior tilting and asymmetric localization of motile primary cilia. Nat. Cell Biol. 12, 407–412.

Cantagrel, V., Silhavy, J. L., Bielas, S. L., Swistun, D., Marsh, S. E., Bertrand, J. Y., Audollent, S., Attié-Bitach, T., Holden, K. R., Dobyns, W. B., et al. (2008). Mutations in the cilia gene ARL13B lead to the classical form of Joubert syndrome. Am. J. Hum. Genet. 83, 170–179.

Caspary, T., Larkins, C. E. and Anderson, K. V. (2007). The graded response to Sonic hedgehog depends on cilia architecture. Dev. Cell 12, 767–778.

Cevik, S., Sanders, A. A. W. M., Van Wijk, E., Boldt, K., Clarke, L., Van Reeuwijk, J., Hori, Y., Horn, N., Hetterschijt, L., Wdowicz, A., et al. (2013). Active transport and diffusion barriers restrict Joubert Syndrome-associated ARL13B/ARL-13 to an Inv-like ciliary membrane subdomain. PLoS Genet. 9, e1003977.

Cho, J. H., Li, Z. A., Zhu, L., Muegge, B. D., Roseman, H. F., Lee, E. Y., Utterback, T., Woodhams, L. G., Bayly, P. V. and Hughes, J. W. (2022). Islet primary cilia motility controls insulin secretion. Sci. Adv. 8, eabq8486.

Davis, R., Swiderski, R., Rahmouni, K., Nishimura, D., Mullins, R., Agassandian, K., Philp, A., Searby, C., Andrews, M., Thompson, S., et al. (2007). A knockin mouse model of the Bardet–Biedl syndrome 1 M390R mutation has cilia defects, ventriculomegaly, retinopathy, and obesity. Proc. Natl. Acad. Sci. U. S. A. 104, 19422–19427.

Delling, M., DeCaen, P. G., Doerner, J. F., Febvay, S. and Clapham, D. E. (2013). Primary cilia are specialized calcium signalling organelles. Nature 504, 311–314.

Dilan, T. L., Moye, A. R., Salido, E. M., Saravanan, T., Kolandaivelu, S., Goldberg, A. F. X. and Ramamurthy, V. (2019). ARL13B, a Joubert Syndrome-associated protein, is critical for retinogenesis and elaboration of mouse photoreceptor outer segments. J. Neurosci. 39, 1347–1364.

Duldulao, N. A., Lee, S. and Sun, Z. (2009). Cilia localization is essential for in vivo functions of the Joubert syndrome protein Arl13b/Scorpion. Development 136, 4033–4042.

Eisa, A., Dey, S., Ignatious, A., Nofal, W., Hess, R. A., Kurokawa, M., Kline, D. and Vijayaraghavan, S. (2021). The protein YWHAE (14-3-3 epsilon) in spermatozoa is essential for male fertility. Andrology 9, 312–328.

Fiore, L., Takata, N., Acosta, S., Ma, W., Pandit, T., Oxendine, M. and Oliver, G. (2020). Optic vesicle morphogenesis requires primary cilia. Dev. Biol. 462, 119–128.

Ford, M. J., Yeyati, P. L., Mali, G. R., Keighren, M. A., Waddell, S. H., Mjoseng, H. K., Douglas, A. T., Hall, E. A., Sakaue-Sawano, A., Miyawaki, A., et al. (2018). A cell/cilia cycle biosensor for single-cell kinetics reveals persistence of cilia after G1/S transition is a general property in cells and mice. Dev. Cell 47, 509–523.e5.

Gigante, E. D., Taylor, M. R., Ivanova, A. A., Kahn, R. A. and Caspary, T. (2020). ARL13B regulates Sonic hedgehog signaling from outside primary cilia. eLife 9, e50434.

Girardet, L., Bernet, A., Calvo, E., Soulet, D., Joly-Beauparlant, C., Droit, A., Cyr, D. G. and Belleannée, C. (2020). Hedgehog signaling pathway regulates gene expression profile of epididymal principal cells through the primary cilium. FASEB J. 34, 7593–7609.

Girardet, L., Cyr, D. G. and Belleannée, C. (2022). Arl13b controls basal cell stemness properties and Hedgehog signaling in the mouse epididymis. Cell Mol. Life Sci. 79, 556.

Grassi, S., Bisconti, M., Martinet, B., Arcolia, V., Simon, J.-F., Wattiez, R., Leroy, B. and Hennebert, E. (2022). Targeted analysis of HSP70 isoforms in human spermatozoa in the context of capacitation and motility. Int. J. Mol. Sci. 23, 6497.

Guo, J., Otis, J. M., Higginbotham, H., Monckton, C., Cheng, J., Asokan, A., Mykytyn, K., Caspary, T., Stuber, G. D. and Anton, E. S. (2017). Primary cilia signaling shapes the development of interneuronal connectivity. Dev. Cell 42, 286–300.e4.

Habif, J. C., Xie, C., De Celis, C., Ukhanov, K., Green, W. W., Moretta, J. C., Zhang, L., Campbell, R. J. and Martens, J. R. (2023). The role of a ciliary GTPase in the regulation of neuronal maturation of olfactory sensory neurons. Development 150, dev201116.

Hanke-Gogokhia, C., Wu, Z., Sharif, A., Yazigi, H., Frederick, J. M. and Baehr, W. (2017). The guanine nucleotide exchange factor Arf-like protein 13b is essential for assembly of the mouse photoreceptor transition zone and outer segment. J. Biol. Chem. 292, 21442–21456.

Hansen, J. N., Rassmann, S., Stüven, B., Jurisch-Yaksi, N. and Wachten, D. (2021). CiliaQ: a simple, open-source software for automated quantification of ciliary morphology and fluorescence in 2D, 3D, and 4D images. Eur. Phys. J. E Soft Matter 44, 18.

Higginbotham, H., Eom, T.-Y., Mariani, L. E., Bachleda, A., Hirt, J., Gukassyan, V., Cusack, C. L., Lai, C., Caspary, T. and Anton, E. S. (2012). Arl13b in primary cilia regulates the migration and placement of interneurons in the developing cerebral cortex. Dev. Cell 23, 925–938.

Higginbotham, H., Guo, J., Yokota, Y., Umberger, N. L., Su, C.-Y., Li, J., Verma, N., Hirt, J., Ghukasyan, V., Caspary, T., et al. (2013). Arl13b-regulated cilia activities are essential for polarized radial glial scaffold formation. Nat. Neurosci. 16, 1000–1007.

Huangfu, D., Liu, A., Rakeman, A. S., Murcia, N. S., Niswander, L. and Anderson, K. V. (2003). Hedgehog signalling in the mouse requires intraflagellar transport proteins. Nature 426, 83–87.

Joiner, A. M., Green, W. W., McIntyre, J. C., Allen, B. L., Schwob, J. E. and Martens, J. R. (2015). Primary cilia on horizontal basal cells regulate regeneration of the olfactory epithelium. J. Neurosci. 35, 13761–13772.

Konno, A., Shiba, K., Cai, C. and Inaba, K. (2015). Branchial cilia and sperm flagella recruit distinct axonemal components. PloS One 10, e0126005.

Larkins, C. E., Aviles, G. D. G., East, M. P., Kahn, R. A. and Caspary, T. (2011). Arl13b regulates ciliogenesis and the dynamic localization of Shh signaling proteins. Mol. Biol. Cell 22, 4694–4703.

Li, Y., Tian, X., Ma, M., Jerman, S., Kong, S., Somlo, S. and Sun, Z. (2016). Deletion of ADP ribosylation factor-like GTPase 13B leads to kidney cysts. J. Am. Soc. Nephrol. 27, 3628–3638.

Li, X., Yang, S., Deepak, V., Chinipardaz, Z. and Yang, S. (2021). Identification of cilia in different mouse tissues. Cells 10, 1623.

Li, Z. A., Cho, J. H., Woodhams, L. G. and Hughes, J. W. (2022). Fluorescence imaging of beta cell primary cilia. Front. Endocrinol. 13, 1004136.

Liu, H., Li, W., Zhang, Y., Zhang, Z., Shang, X., Zhang, L., Zhang, S., Li, Y., Somoza, A. V., Delpi, B., et al. (2017). IFT25, an intraflagellar transporter protein dispensable for ciliogenesis in somatic cells, is essential for sperm flagella formation. Biol. Reprod. 96, 993–1006.

Liu, Y.-X., Li, W.-J., Zhang, R.-K., Sun, S.-N. and Fan, Z.-C. (2023). Unraveling the intricate cargo-BBSome coupling mechanism at the ciliary tip. Proc. Natl. Acad. Sci. 120, e2218819120.

Lu, H., Toh, M. T., Narasimhan, V., Thamilselvam, S. K., Choksi, S. P. and Roy, S. (2015). A function for the Joubert syndrome protein Arl13b in ciliary membrane extension and ciliary length regulation. Dev. Biol. 397, 225–236.

Mariani, L. E., Bijlsma, M. F., Ivanova, A. A., Suciu, S. K., Kahn, R. A. and Caspary, T. (2016). Arl13b regulates Shh signaling from both inside and outside the cilium. Mol. Biol. Cell 27, 3780–3790.

Mykytyn, K., Mullins, R. F., Andrews, M., Chiang, A. P., Swiderski, R. E., Yang, B., Braun, T., Casavant, T., Stone, E. M. and Sheffield, V. C. (2004). Bardet–Biedl syndrome type 4 (BBS4)-null mice implicate Bbs4 in flagella formation but not global cilia assembly. Proc. Natl. Acad. Sci. 101, 8664–8669.

Nielsen, M. G. and Raff, E. C. (2002). The best of all worlds or the best possible world? Developmental constraint in the evolution of β-tubulin and the sperm tail axoneme. Evol. Dev. 4, 303–315.

Nishimura, D. Y., Fath, M., Mullins, R. F., Searby, C., Andrews, M., Davis, R., Andorf, J. L., Mykytyn, K., Swiderski, R. E., Yang, B., et al. (2004). Bbs2-null mice have neurosensory deficits, a defect in social dominance, and retinopathy associated with mislocalization of rhodopsin. Proc. Natl. Acad. Sci. 101, 16588–16593.

Nozaki, S., Katoh, Y., Terada, M., Michisaka, S., Funabashi, T., Takahashi, S., Kontani, K. and Nakayama, K. (2017). Regulation of ciliary retrograde protein trafficking by the Joubert syndrome proteins ARL13B and INPP5E. J. Cell Sci. 130, 563–576.

Ocbina, P. J. R., Eggenschwiler, J. T., Moskowitz, I. and Anderson, K. V. (2011). Complex interactions between genes controlling trafficking in primary cilia. Nat. Genet. 43, 547–553.

Osawa, Y., Murata, K., Usui, M., Kuba, Y., Le, H. T., Mikami, N., Nakagawa, T., Daitoku, Y., Kato, K., Shawki, H. H., et al. (2021). EXOC1 plays an integral role in spermatogonia pseudopod elongation and spermatocyte stable syncytium formation in mice. eLife 10, e59759.

Pintado, P., Sampaio, P., Tavares, B., Montenegro-Johnson, T. D., Smith, D. J. and Lopes, S. S. (2017). Dynamics of cilia length in left-right development. R. Soc. Open Sci. 4, 161102.

Sakaue-Sawano, A., Kurokawa, H., Morimura, T., Hanyu, A., Hama, H., Osawa, H., Kashiwagi, S., Fukami, K., Miyata, T., Miyoshi, H., et al. (2008). Visualizing spatiotemporal dynamics of multicellular cell-cycle progression. Cell 132, 487–498.

San Agustin, J. T., Pazour, G. J. and Witman, G. B. (2015). Intraflagellar transport is essential for mammalian spermiogenesis but is absent in mature sperm. Mol. Biol. Cell 26, 4358–4372.

Schindelin, J., Arganda-Carreras, I., Frise, E., Kaynig, V., Longair, M., Pietzsch, T., Preibisch, S., Rueden, C., Saalfeld, S., Schmid, B., et al. (2012). Fiji: an open-source platform for biological-image analysis. Nat. Methods 9, 676–682.

Schmitz, F., Burtscher, I., Stauber, M., Gossler, A. and Lickert, H. (2017). A novel Cre-inducible knock-in ARL13B-tRFP fusion cilium reporter. Genesis 55, e23073.

Seixas, C., Choi, S. Y., Polgar, N., Umberger, N. L., East, M. P., Zuo, X., Moreiras, H., Ghossoub, R., Benmerah, A., Kahn, R. A., et al. (2016). Arl13b and the exocyst interact synergistically in ciliogenesis. Mol. Biol. Cell 27, 308–320.

Shi, L., Zhou, T., Huang, Q., Zhang, S., Li, W., Zhang, L., Hess, R. A., Pazour, G. J. and Zhang, Z. (2019). Intraflagellar transport protein 74 is essential for spermatogenesis and male fertility in mice. Biol. Reprod. 101, 188–199.

Su, C.-Y., Bay, S. N., Mariani, L. E., Hillman, M. J. and Caspary, T. (2012). Temporal deletion of *Arl13b* reveals that a mispatterned neural tube corrects cell fate over time. Development 139, 4062–4071.

Suciu, S. K., Long, A. B. and Caspary, T. (2021). Smoothened and ARL13B are critical in mouse for superior cerebellar peduncle targeting. Genetics 218, iyab084.

Sun, Z., Amsterdam, A., Pazour, G. J., Cole, D. G., Miller, M. S. and Hopkins, N. (2004). A genetic screen in zebrafish identifies cilia genes as a principal cause of cystic kidney. Development 131, 4085–4093.

Terry, T. T., Gigante, E. D., Alexandre, C. M., Brewer, K. M., Yue, X., Berbari, N. F., Vaisse, C. and Caspary, T. (2025). Ciliary ARL13B is essential for body weight regulation in adult mice. bioRxiv doi:10.1101/2023.08.02.551695.

Thomas, S., Cantagrel, V., Mariani, L., Serre, V., Lee, J.-E., Elkhartoufi, N., De Lonlay, P., Desguerre, I., Munnich, A., Boddaert, N., et al. (2015). Identification of a novel ARL13B variant in a Joubert syndrome-affected patient with retinal impairment and obesity. Eur. J. Hum. Genet. 23, 621–627.

Van Sciver, R. E., Long, A. B., Katz, H. G., Gigante, E. D. and Caspary, T. (2023). Ciliary ARL13B inhibits developmental kidney cystogenesis in mouse. Dev. Biol. 500, 1–9.

Zhang, Q., Li, Y., Zhang, Y., Torres, V. E., Harris, P. C., Ling, K. and Hu, J. (2016). GTP-binding of ARL-3 is activated by ARL-13 as a GEF and stabilized by UNC-119. Sci. Rep. 6, 24534.

Zhang, Y., Liu, H., Li, W., Zhang, Z., Shang, X., Zhang, D., Li, Y., Zhang, S., Liu, J., Hess, R. A., et al. (2017). Intraflagellar transporter protein (IFT27), an IFT25 binding partner, is essential for male fertility and spermiogenesis in mice. Dev. Biol. 432, 125–139.

