## Supplementary figures and images for "ARL13B-Cerulean rescues *Arl13b*-null mouse from embryonic lethality and reveals a role for ARL13B in spermatogenesis"

### Supplemental Figures

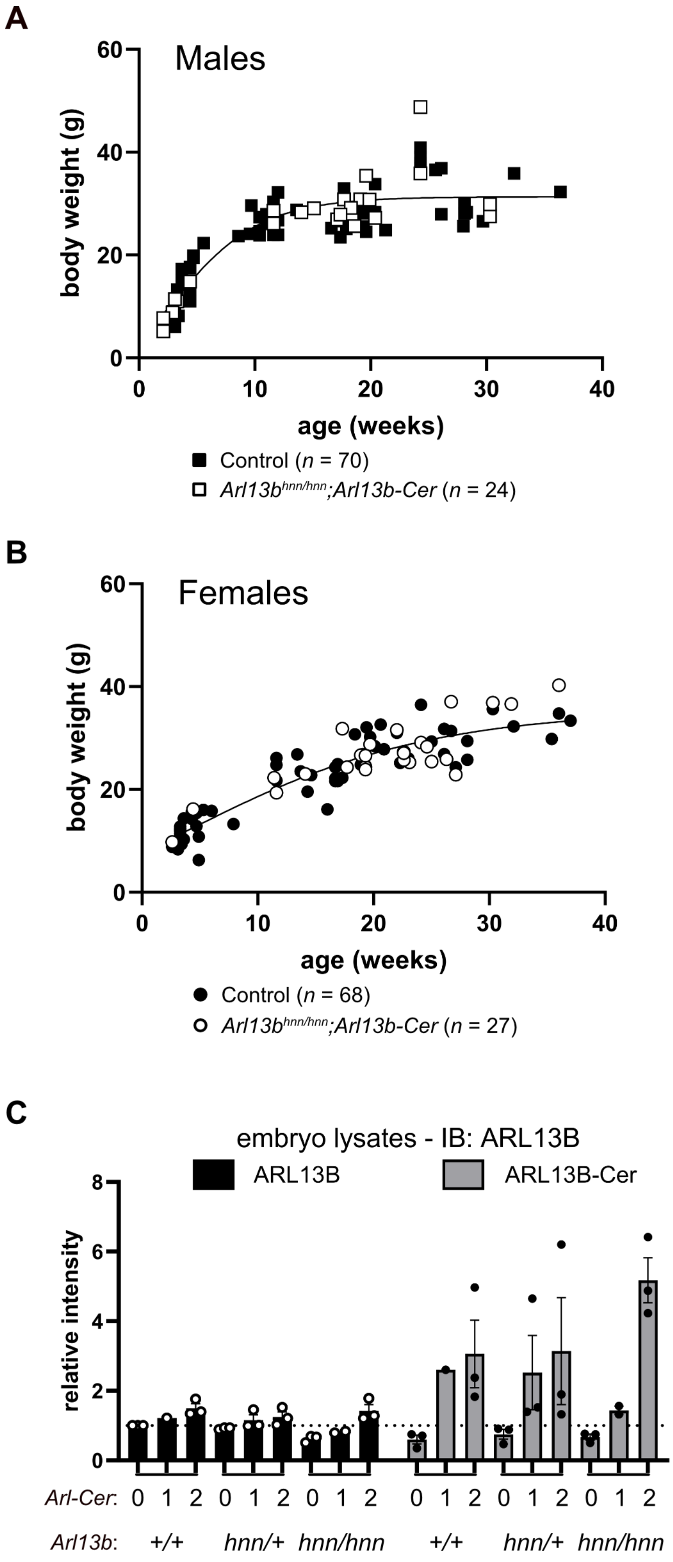

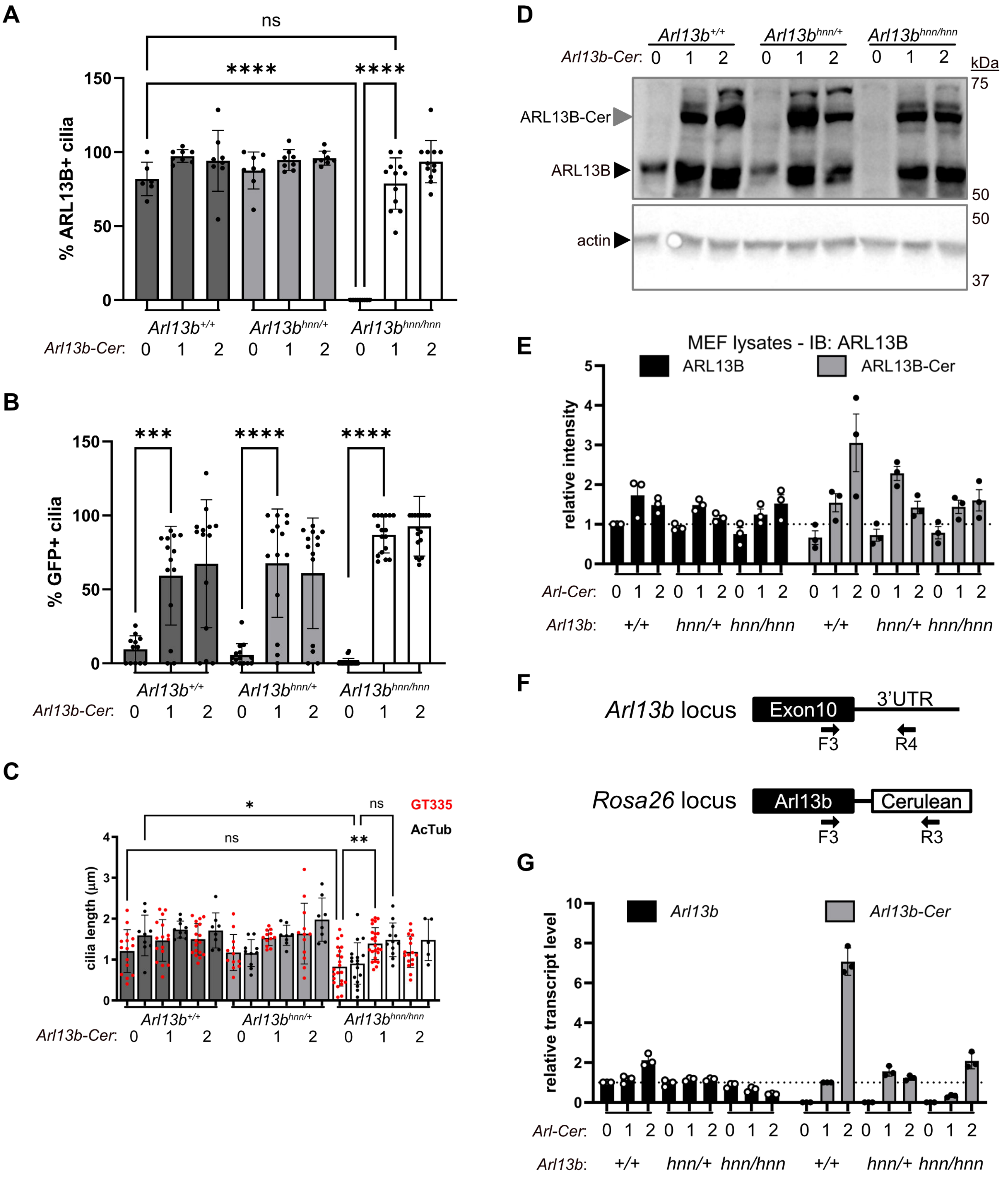

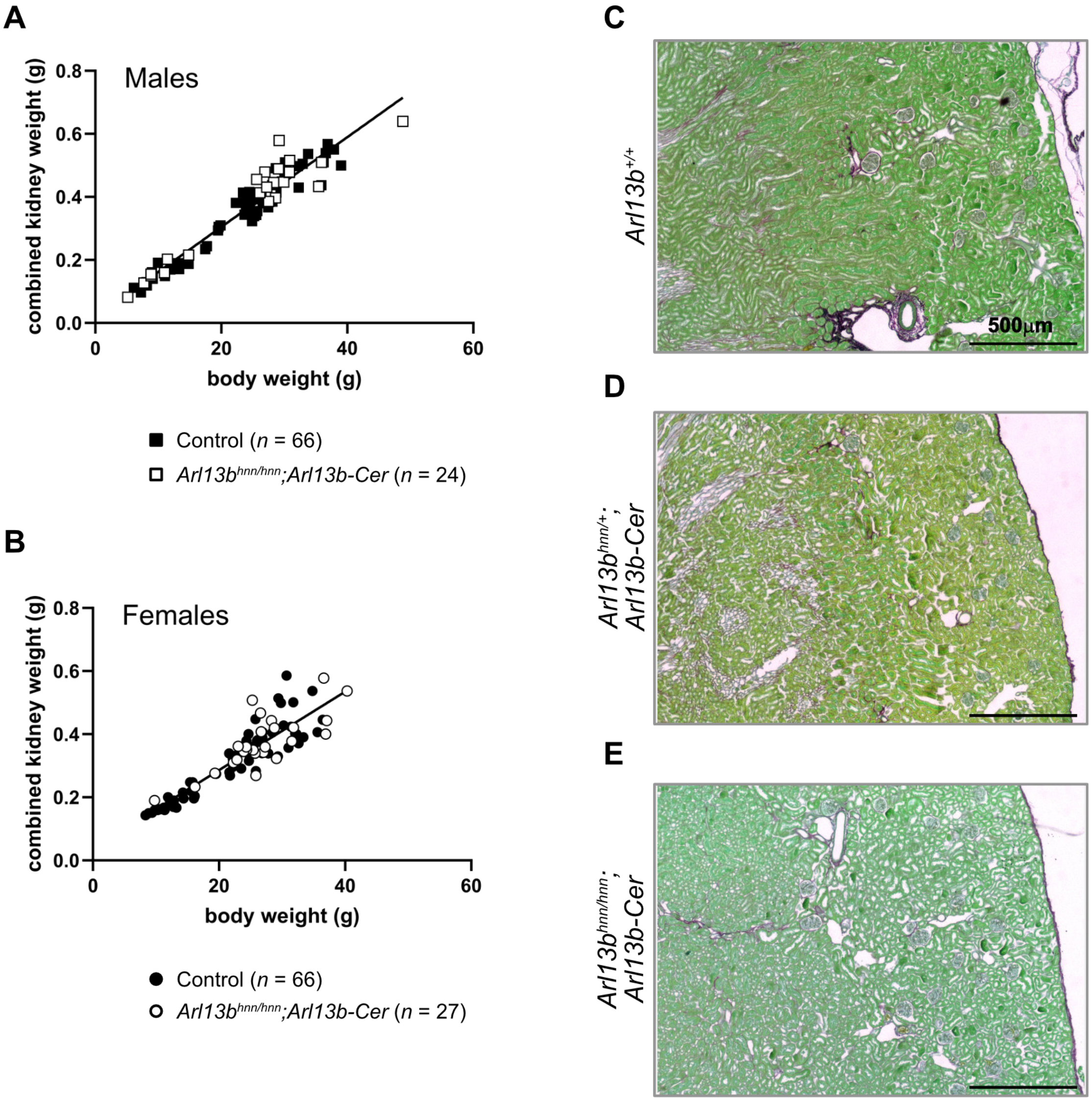

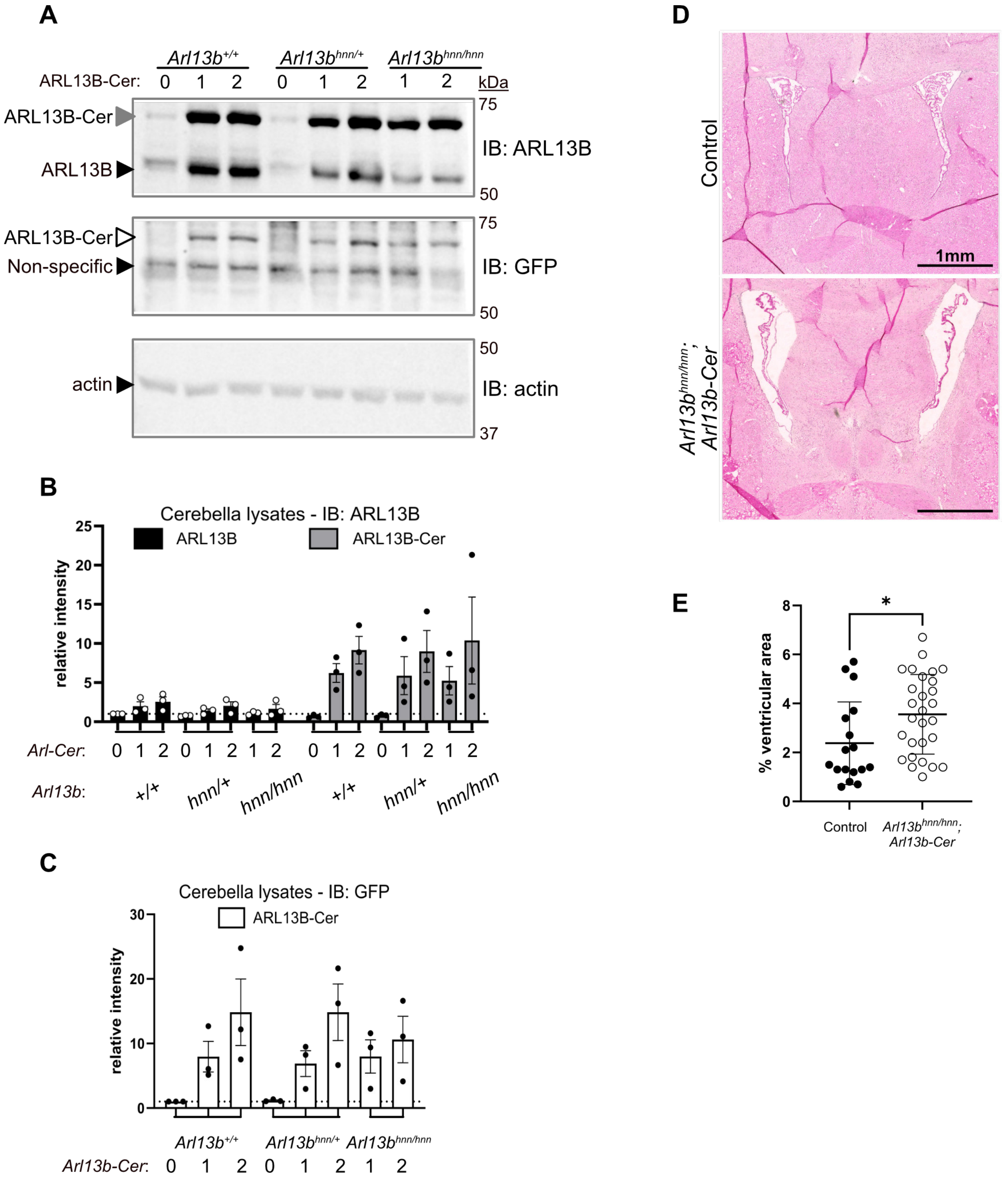

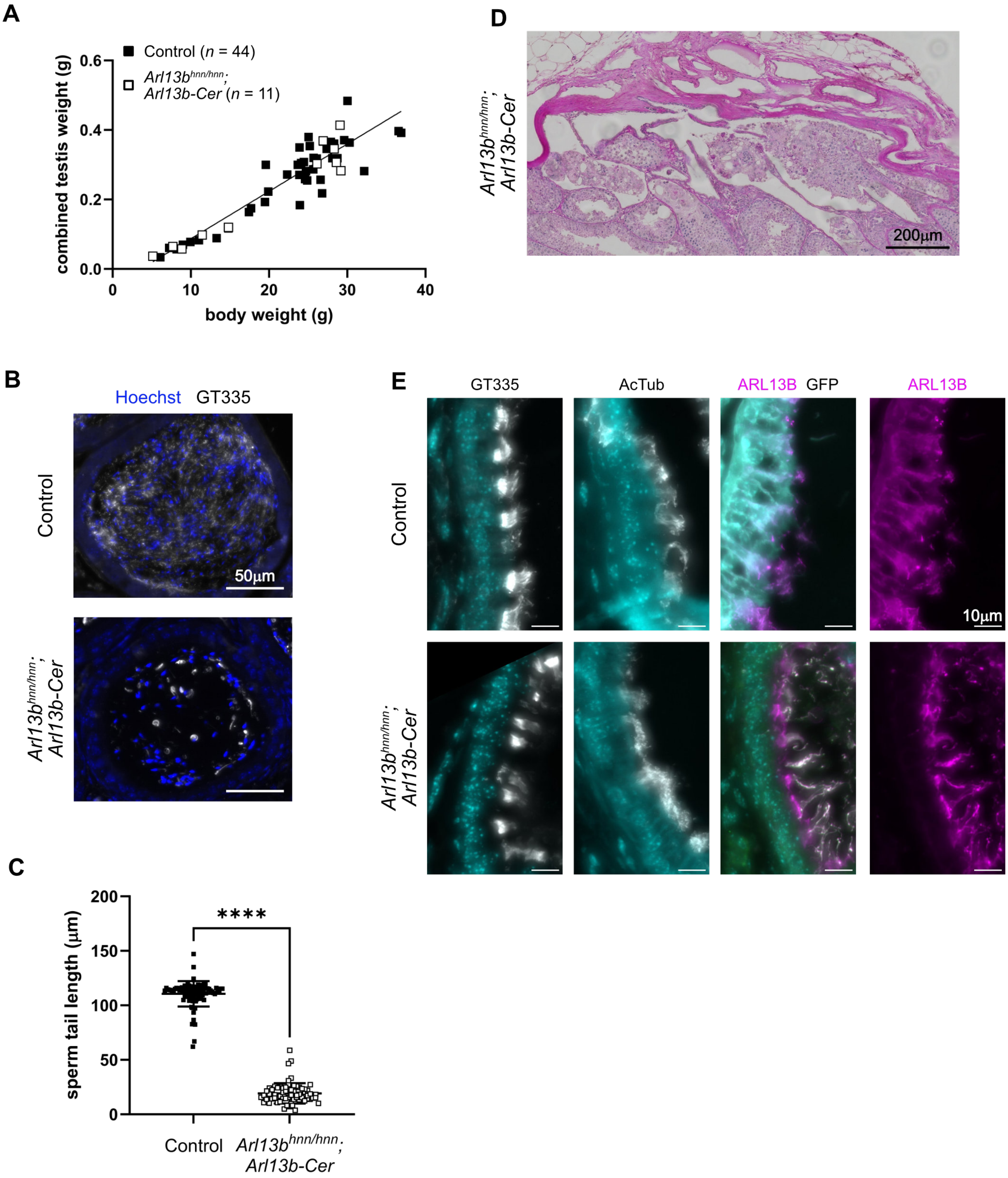

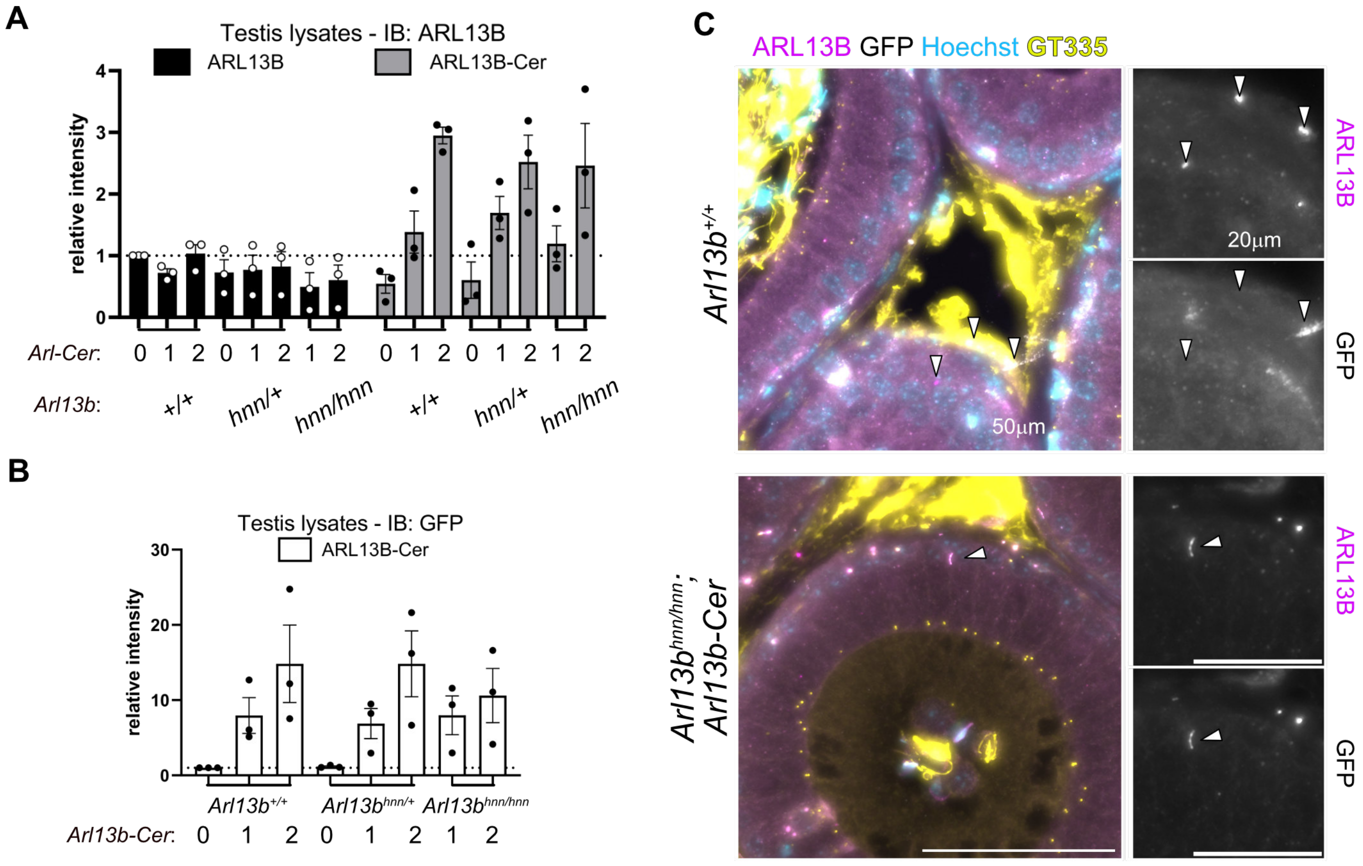
